## Supplementary material for "Chloroplast acquisition without the gene transfer in kleptoplastic sea slugs, *Plakobranchus ocellatus*": Supple_file_10.docx

Enriched GOs of the significantly upregulated genes on the digestive gland of *P. ocellatus*

| GO term description | Over represented p-value | DEG number from Poc | Total gene number in Poc |  | DG-upregulated gene belong the GO |
| --- | --- | --- | --- | --- | --- |
| GO:0004190 aspartic-type endopeptidase activity | 2.56E-71 | 45 | 288 |  | p1440c63.113, p664c69.11, p855c67.174, p807c65.17, p1440c63.126, p3924c64.18, p1440c63.80, p543c70.163, p1440c63.23, p1440c63.36, p1440c63.12, p3924c64.17, p1440c63.63, p775c64.19, p3924c64.1, p665c65.51, p664c69.19, p428c65.20, p1440c63.105, p665c65.34, p445c59.46, p1440c63.3, p1412c61.10, p1440c63.11, p1440c63.8, p855c67.163, p1440c63.84, p1015c69.247, p665c65.52, p596c69.2, p329c71.312, p855c67.168, p775c64.23, p2791c66.18, p1440c63.43, p3924c64.35, p238c64.67, p445c59.50, p8162c67.79, p737c53.70, p3924c64.16, p775c64.20, p1440c63.115, p1440c63.119, p665c65.58, |
| GO:0006508 proteolysis | 1.65E-60 | 49 | 684 |  | p1440c63.113, p664c69.11, p855c67.174, p807c65.17, p1440c63.126, p3924c64.18, p1440c63.80, p543c70.163, p1440c63.23, p1440c63.36, p1440c63.12, p3924c64.17, p1440c63.63, p775c64.19, p3924c64.1, p665c65.51, p664c69.19, p428c65.20, p1440c63.105, p665c65.34, p445c59.46, p1440c63.3, p1412c61.10, p1440c63.11, p1440c63.8, p855c67.163, p1440c63.84, p1015c69.247, p665c65.52, p596c69.2, p329c71.312, p855c67.168, p775c64.23, p2791c66.18, p1440c63.43, p3924c64.35, p238c64.67, p445c59.50, p8162c67.79, p737c53.70, p3924c64.16, p775c64.20, p1440c63.115, p1440c63.119, p1510c68.4, p89c59.118, p22c64.145, p665c65.58, p258731c69.52, |
| GO:0004869 cysteine-type endopeptidase inhibitor activity | 6.52E-05 | 2 | 6 |  | p34c66.53, p34c66.54, |
| GO:0006952 defense response | 1.95E-04 | 2 | 10 |  | p4035c66.3, p606c64.1, |
| GO:0043169 cation binding | 2.38E-04 | 2 | 11 |  | p128c64.183, p1548c63.54, |
| GO:0006885 regulation of pH | 3.36E-04 | 2 | 13 |  | p11c67.36, p1947c65.69, |
| GO:0006730 one-carbon metabolic process | 4.51E-04 | 2 | 15 |  | p1053c67.29, p1947c65.69, |
| GO:0005975 carbohydrate metabolic process | 4.67E-04 | 4 | 169 |  | p590c70.27, p128c64.183, p105c62.18, p1548c63.54, |
| GO:0048856 anatomical structure development | 9.83E-04 | 2 | 22 |  | p1440c63.115, p1440c63.119, |
