## Supplementary material for "Chloroplast acquisition without the gene transfer in kleptoplastic sea slugs, *Plakobranchus ocellatus*": Supple_file_15.docx

Result of OrthoFinder analysis with additional data from *E. chlorotica* gene set (gene counts)

| Group name | Ema | PoB | Ecl | Api | Aca | Bfl | Bgl | Bmo | Cte | Cgi | Dme | Hdi | Lan | Lgi | Mus | Obi | Dre | Has | Pfu |
| --- | --- | --- | --- | --- | --- | --- | --- | --- | --- | --- | --- | --- | --- | --- | --- | --- | --- | --- | --- |
| Cathepsin D-like | 5**^A^** | 203**^B^** | 4**^C^** | 3 | 5 | 5 | 4 | 3 | 1 | 2 | 19 | 0 | 3 | 3 | 7 | 2 | 17 | 8 | 1 |
| Lectin-like | 214**^D^** | 410**^E^** | 72**^F^** | 3 | 46 | 398 | 126 | 11 | 128 | 275 | 28 | 96 | 133 | 62 | 44 | 10 | 105 | 37 | 166 |
| Apolipoprotein D-like | 8**^G^** | 36**^H^** | 3**^I^** | 85 | 1 | 0 | 0 | 14 | 11 | 54 | 7 | 22 | 8 | 15 | 4 | 5 | 8 | 4 | 1 |

**A:** e2244c39.16, e2244c39.20, e4981c37.4, e4981c37.5, e8012c40.2

**B:** p1015c69.245, p1015c69.247, p1106c69.41, p1106c69.44, p1412c61.10, p1412c61.2, p1440c63.10, p1440c63.100, p1440c63.101, p1440c63.102, p1440c63.103, p1440c63.104, p1440c63.105, p1440c63.106, p1440c63.11, p1440c63.113, p1440c63.115, p1440c63.117, p1440c63.119, p1440c63.12, p1440c63.120, p1440c63.122, p1440c63.124, p1440c63.125, p1440c63.126, p1440c63.127, p1440c63.14, p1440c63.15, p1440c63.2, p1440c63.20, p1440c63.23, p1440c63.26, p1440c63.27, p1440c63.28, p1440c63.3, p1440c63.30, p1440c63.36, p1440c63.42, p1440c63.43, p1440c63.44, p1440c63.45, p1440c63.51, p1440c63.52, p1440c63.55, p1440c63.58, p1440c63.63, p1440c63.69, p1440c63.7, p1440c63.73, p1440c63.76, p1440c63.77, p1440c63.79, p1440c63.8, p1440c63.80, p1440c63.81, p1440c63.83, p1440c63.84, p1440c63.89, p1440c63.9, p1440c63.90, p1440c63.91, p1440c63.96, p1440c63.98, p1525c58.1, p1529c64.14, p17174c35.1, p183713c35.1, p1860c62.49, p1860c62.50, p1860c62.51, p19234c41.1, p2015c61.59, p2057c61.46, p2057c61.51, p216c65.11, p216c65.8, p22c64.171, p238c64.56, p238c64.58, p238c64.60, p238c64.61, p238c64.67, p254c66.117, p255c64.80, p2574c60.31, p2574c60.32, p2574c60.35, p258745c64.77, p266c63.161, p2791c66.17, p2791c66.18, p30156c39.1, p30376c56.1, p30807c69.1, p329c71.310, p329c71.312, p333c67.117, p333c67.123, p333c67.126, p333c67.128, p34996c59.1, p35389c41.1, p3560c63.43, p35754c38.1, p3605c51.5, p374c67.44, p374c67.53, p3924c64.1, p3924c64.14, p3924c64.16, p3924c64.17, p3924c64.18, p3924c64.19, p3924c64.21, p3924c64.32, p3924c64.35, p3924c64.4, p3924c64.5, p3924c64.6, p3924c64.78, p428c65.20, p445c59.36, p445c59.38, p445c59.43, p445c59.46, p445c59.50, p445c59.54, p445c59.57, p445c59.58, p445c59.61, p445c59.62, p445c59.64, p445c59.66, p51766c46.3, p543c70.163, p56091c35.1, p596c69.2, p59943c48.1, p609c69.52, p609c69.84, p63233c38.1, p664c69.11, p664c69.13, p664c69.15, p664c69.19, p664c69.24, p664c69.25, p664c69.32, p664c69.34, p665c65.34, p665c65.40, p665c65.47, p665c65.48, p665c65.49, p665c65.50, p665c65.51, p665c65.52, p665c65.53, p665c65.57, p665c65.58, p665c65.59, p665c65.62, p665c65.65, p737c53.70, p737c53.71, p737c53.72, p775c64.1, p775c64.15, p775c64.16, p775c64.19, p775c64.2, p775c64.20, p775c64.23, p800c68.116, p807c65.16, p807c65.17, p807c65.22, p807c65.25, p807c65.27, p807c65.38, p807c65.41, p807c65.42, p8162c67.103, p8162c67.108, p8162c67.48, p8162c67.79, p8162c67.82, p8162c67.88, p8162c67.89, p8491c59.1, p8558c44.1, p855c67.161, p855c67.162, p855c67.163, p855c67.164, p855c67.168, p855c67.171, p855c67.172, p855c67.174, p864c38.3, p93400c36.1, p947c59.272, p97c59.82

**C:**^,^RUS74941.1, RUS87636.1, RUS87637.1, RUS89981.1

**D:** e10094c43.1, e10094c43.2, e101c35.8, e10327c38.3, e103301c44.16, e10462c41.1, e10655c37.1, e10666c38.1, e10764c35.2, e10764c35.3, e1113c45.25, e1116c29.10, e1121c45.40, e1131c43.70, e11361c34.21, e11595c36.1, e1164c44.4, e1164c44.9, e11668c37.2, e11880c35.1, e11969c36.4, e12295c25.4, e12486c38.7, e12518c40.3, e12518c40.5, e12595c42.5, e12595c42.6, e12595c42.7, e12651c38.5, e12682c48.9, e1275c33.6, e128757c48.1, e13123c30.5, e1318c44.48, e13376c36.12, e14309c38.2, e14600c41.2, e1496c33.3, e1516c38.8, e15522c35.1, e15573c33.1, e15859c36.9, e1598c45.65, e16001c30.2, e16112c35.14, e1652c45.28, e1653c43.13, e16666c32.1, e16666c32.2, e16666c32.5, e16777c31.1, e17217c35.4, e17234c30.2, e1732c40.7, e1733c40.17, e174475c32.1, e1770c40.1, e18174c38.12, e18174c38.9, e1895c40.21, e19587c27.2, e19625c44.21, e19625c44.24, e1967c35.2, e1967c35.4, e1994c41.12, e20336c40.1, e20336c40.2, e210547c19.2, e212284c41.7, e212290c37.5, e212290c37.7, e2125c39.2, e2125c39.5, e214759c45.7, e216082c42.8, e2177c43.3, e224005c42.2, e224005c42.3, e224005c42.4, e224099c35.1, e22489c36.2, e2330c38.9, e23356c42.1, e2366c41.32, e2366c41.33, e2366c41.34, e2391c33.8, e24512c39.1, e2538c33.7, e2632c42.1, e2640c39.3, e27845c35.3, e280c43.52, e280c43.54, e28258c37.9, e2891c46.44, e2924c47.1, e2936c41.16, e2951c42.3, e3030c36.12, e30461c43.16, e308c40.33, e3309c35.4, e3349c37.4, e3489c45.48, e3573c41.1, e3702c38.31, e3733c45.9, e374c43.6, e3770c38.30, e37732c19.1, e379c31.2, e3909c35.9, e3950c37.4, e3991c44.11, e4013c42.2, e4139c37.5, e413c40.1, e4195c32.3, e4278c41.3, e4409c43.1, e4409c43.5, e4497c33.9, e4517c36.4, e4572c33.2, e463c36.10, e46448c25.1, e4674c39.16, e4674c39.27, e46971c29.1, e4740c32.8, e47886c33.1, e4788c40.4, e4810c33.3, e4897c45.38, e499c42.5, e502c47.6, e502c47.7, e5202c34.8, e5323c35.2, e54608c41.1, e5559c41.2, e5626c39.2, e5661c36.2, e5716c37.7, e5744c33.1, e5786c35.2, e5869c35.14, e588c41.23, e588c41.27, e588c41.30, e588c41.33, e588c41.35, e5c41.4, e6009c34.4, e603c38.41, e6182c43.13, e6444c46.9, e6489c44.4, e6611c34.4, e6621c41.8, e6631c31.3, e66797c25.1, e6837c39.10, e6837c39.8, e6863c45.35, e6889c38.3, e6922c33.3, e6924c35.6, e6926c34.11, e6926c34.3, e6926c34.5, e7085c37.9, e7329c41.8, e7333c41.4, e7372c37.4, e7390c42.26, e7390c42.30, e7487c37.10, e7487c37.13, e7487c37.8, e7488c35.1, e75968c25.1, e76442c41.1, e7786c30.1, e7837c39.12, e7856c39.22, e785c40.4, e7944c38.7, e8025c35.21, e8068c38.1, e8350c35.4, e83820c30.1, e8532c37.3, e8598c31.1, e8728c35.1, e8901c40.1, e8901c40.4, e9032c37.14, e934c37.40, e946c31.2, e954c34.13, e954c34.33, e955c43.19, e955c43.22, e955c43.23, e955c43.24, e955c43.25, e955c43.27, e955c43.29, e955c43.30, e955c43.31, e9716c25.5

**E:** p1013c63.45, p1015c69.241, p1015c69.5, p10278c41.2, p1039c62.2, p108080c46.1, p110c62.105, p1121c63.100, p1121c63.130, p1121c63.97, p1121c63.98, p1147c61.57, p1147c61.59, p114c62.106, p114c62.114, p114c62.97, p114c62.99, p115899c41.1, p118c63.26, p118c63.39, p118c63.42, p118c63.50, p118c63.51, p118c63.59, p118c63.61, p118c63.73, p1194c63.105, p1209c50.1, p1215c59.122, p12237c46.1, p12294c64.10, p123c64.16, p124c71.257, p124c71.259, p124c71.261, p124c71.262, p124c71.264, p124c71.266, p12763c48.1, p1321c62.95, p1352c59.54, p14216c54.2, p1425c64.196, p1425c64.197, p1425c64.198, p1425c64.92, p1427c68.91, p143c69.9, p145c61.17, p1464c63.83, p1511c63.56, p1511c63.58, p1516c67.88, p1529c64.11, p1529c64.37, p1531c64.13, p1531c64.9, p1581c68.1, p1581c68.2, p16742c40.7, p1686c67.2, p1686c67.3, p16c66.82, p170c62.58, p170c62.59, p170c62.62, p171c61.137, p171c61.192, p171c61.196, p1739c63.85, p1739c63.86, p174c61.58, p1753c40.21, p1819c60.9, p1846c62.20, p1857c64.5, p1866c63.111, p1874c66.27, p1890c66.15, p1941c66.1, p1949c65.33, p1959c71.21, p1959c71.22, p195c62.26, p195c62.27, p1987c67.33, p2021c67.18, p2051c59.50, p2053c66.138, p2092c44.45, p2102c68.10, p2102c68.17, p2114c60.7, p2150c71.1, p2195c58.10, p2195c58.11, p2195c58.2, p2195c58.23, p2195c58.4, p2195c58.5, p2197c68.142, p2251c65.17, p2251c65.52, p226c65.101, p227c62.10, p227c62.128, p227c62.166, p227c62.168, p227c62.170, p227c62.191, p227c62.197, p227c62.26, p227c62.27, p227c62.29, p227c62.33, p227c62.36, p227c62.37, p227c62.38, p227c62.56, p227c62.6, p227c62.8, p227c62.87, p22c64.198, p23437c72.2, p23437c72.3, p238c64.110, p238c64.116, p2413c58.68, p2437c66.8, p24778c83.1, p24c58.17, p2517c51.4, p2517c51.5, p2530c72.1, p2530c72.90, p255984c65.78, p255c64.82, p256948c53.1, p256c68.13, p256c68.16, p256c68.17, p256c68.23, p256c68.31, p256c68.32, p257049c72.18, p257c63.203, p258731c69.113, p258734c62.168, p258734c62.231, p258734c62.9, p258746c66.13, p258746c66.239, p258746c66.240, p258746c66.257, p258746c66.261, p258746c66.63, p258746c66.70, p258747c66.2, p258747c66.6, p258748c66.169, p258748c66.170, p258748c66.175, p258748c66.178, p258753c65.148, p258753c65.149, p258753c65.173, p258754c68.1, p258754c68.2, p258754c68.4, p258781c59.14, p258781c59.16, p258788c60.13, p258789c70.10, p258789c70.12, p258789c70.8, p258790c64.57, p258822c43.2, p258823c64.1, p258828c43.1, p258836c59.11, p258836c59.6, p258838c66.2, p2634c44.1, p266c63.16, p266c63.18, p266c63.351, p266c63.353, p2701c65.6, p275c64.241, p2994c64.4, p2994c64.5, p3054c66.143, p3054c66.145, p3054c66.150, p3054c66.152, p3054c66.153, p3054c66.73, p3054c66.74, p3054c66.80, p3054c66.81, p3054c66.84, p3054c66.86, p3054c66.90, p311c62.66, p311c62.9, p3159c54.28, p319c70.179, p3219c67.21, p3219c67.22, p3238c66.28, p3265c61.1, p330c75.111, p3334c67.100, p3334c67.101, p3334c67.102, p3334c67.69, p3334c67.75, p3334c67.97, p3334c67.98, p3334c67.99, p335c60.149, p335c60.16, p335c60.22, p335c60.25, p335c60.28, p335c60.39, p335c60.5, p3384c60.111, p3385c70.37, p3385c70.59, p3385c70.7, p3385c70.8, p3388c67.23, p3391c70.39, p339c68.162, p339c68.163, p339c68.166, p339c68.167, p339c68.52, p339c68.64, p339c68.72, p3419c53.2, p3560c63.28, p363c79.86, p37404c69.64, p374c67.34, p374c67.57, p3752c58.2, p3752c58.3, p3752c58.4, p3752c58.5, p3752c58.7, p38383c53.1, p385c65.53, p385c65.54, p3952c56.14, p39968c42.1, p4016c57.48, p401c55.52, p401c55.53, p401c55.54, p415c66.5, p415c66.6, p4212c67.5, p422c65.69, p422c65.70, p422c65.79, p422c65.86, p429c63.241, p4334c64.1, p4334c64.15, p4334c64.20, p4334c64.26, p4334c64.28, p4334c64.3, p4334c64.30, p4334c64.4, p4334c64.44, p4426c53.7, p445c59.2, p459c71.116, p469c60.20, p470c60.18, p473c72.115, p4763c78.1, p47c72.168, p47c72.174, p4867c67.45, p4867c67.46, p4875c60.1, p4875c60.3, p4875c60.7, p48c60.10, p48c60.7, p48c60.92, p494c61.125, p494c61.131, p494c61.132, p494c61.135, p494c61.136, p494c61.141, p494c61.146, p494c61.148, p494c61.150, p494c61.153, p501c63.110, p501c63.48, p501c63.49, p501c63.50, p501c63.56, p501c63.58, p501c63.61, p501c63.63, p501c63.64, p501c63.65, p501c63.67, p515c60.323, p515c60.330, p51c65.70, p51c65.71, p523c60.2, p523c60.3, p523c60.31, p523c60.35, p5367c63.11, p543c70.2, p543c70.3, p543c70.5, p543c70.8, p543c70.93, p560c64.1, p560c64.2, p560c64.3, p560c64.4, p560c64.6, p561c61.5, p590c70.18, p600c62.26, p600c62.88, p600c62.90, p600c62.94, p600c62.95, p600c62.96, p606c64.40, p609c69.2, p609c69.64, p609c69.70, p609c69.8, p609c69.83, p6313c71.6, p6320c73.1, p6356c46.2, p6356c46.3, p639c70.164, p639c70.6, p6430c70.3, p6430c70.6, p650c70.3, p65450c38.1, p68146c61.1, p685c68.151, p6955c62.1, p69c69.67, p6c66.67, p6c66.68, p702c64.15, p702c64.17, p7066c37.1, p706c62.80, p7096c40.3, p7107c59.7, p71441c62.1, p7187c46.2, p7262c64.1, p758c62.46, p758c62.70, p75c56.59, p768c75.4, p776c57.1, p7806c45.1, p7c70.82, p800c68.44, p812c68.55, p813c67.23, p813c67.31, p8162c67.85, p819c55.16, p819c55.4, p819c55.6, p8329c60.25, p8329c60.27, p854c69.60, p855c67.129, p855c67.134, p855c67.208, p8760c46.6, p8760c46.8, p88033c67.1, p883c68.155, p884c71.27, p887c55.25, p89185c30.1, p90525c94.1, p918c66.32, p918c66.33, p92c69.41, p94c72.22, p958c60.20, p958c60.24, p958c60.27

**F:** RUS68616.1, RUS68633.1, RUS68647.1, RUS68662.1, RUS69163.1, RUS69541.1, RUS69648.1, RUS69842.1, RUS70124.1, RUS70475.1, RUS70726.1, RUS71236.1, RUS71340.1, RUS71551.1, RUS72059.1, RUS72139.1, RUS72211.1, RUS73268.1, RUS73297.1, RUS74009.1, RUS74083.1, RUS75111.1, RUS75504.1, RUS75505.1, RUS76015.1, RUS76722.1, RUS76779.1, RUS77171.1, RUS77679.1, RUS80279.1, RUS80935.1, RUS81139.1, RUS81800.1, RUS82192.1, RUS82307.1, RUS82887.1, RUS83253.1, RUS83536.1, RUS83755.1, RUS83907.1, RUS83908.1, RUS84090.1, RUS84091.1, RUS84408.1, RUS84451.1, RUS85257.1, RUS85260.1, RUS85335.1, RUS86035.1, RUS86039.1, RUS86136.1, RUS86160.1, RUS86267.1, RUS86568.1, RUS86702.1, RUS87571.1, RUS87572.1, RUS87614.1, RUS87724.1, RUS87725.1, RUS87726.1, RUS88118.1, RUS88255.1, RUS88257.1, RUS88258.1, RUS88638.1, RUS88639.1, RUS89920.1, RUS90178.1, RUS90265.1, RUS90584.1, RUS91679.1

**G:** e12398c37.5, e15279c22.1, e19340c38.16, e2950c42.1, e358c35.6, e358c35.7, e42367c51.1, e4695c40.1

**H:** p1001c63.5, p11751c81.2, p12498c72.2, p126c77.126, p126c77.67, p1425c64.85, p1662c68.129, p17146c62.2, p189c72.53, p189c72.54, p2081c70.47, p221c62.44, p238c64.62, p2538c67.15, p2550c65.67, p258670c68.31, p258755c69.28, p282c66.68, p329c71.111, p339c68.7, p385c65.19, p38c58.15, p40759c49.1, p4154c75.33, p440c72.42, p473c72.110, p474c68.27, p474c68.28, p509c73.151, p589c77.14, p59c63.60, p6257c70.26, p6430c70.39, p735c63.50, p754c67.20, p838c70.295

I RUS69394.1, RUS74332.1, RUS74836.1
