## Supplementary Figures for "Chloroplast acquisition without the gene transfer in kleptoplastic sea slugs, *Plakobranchus ocellatus*": Figure_4_supple_6.pdf

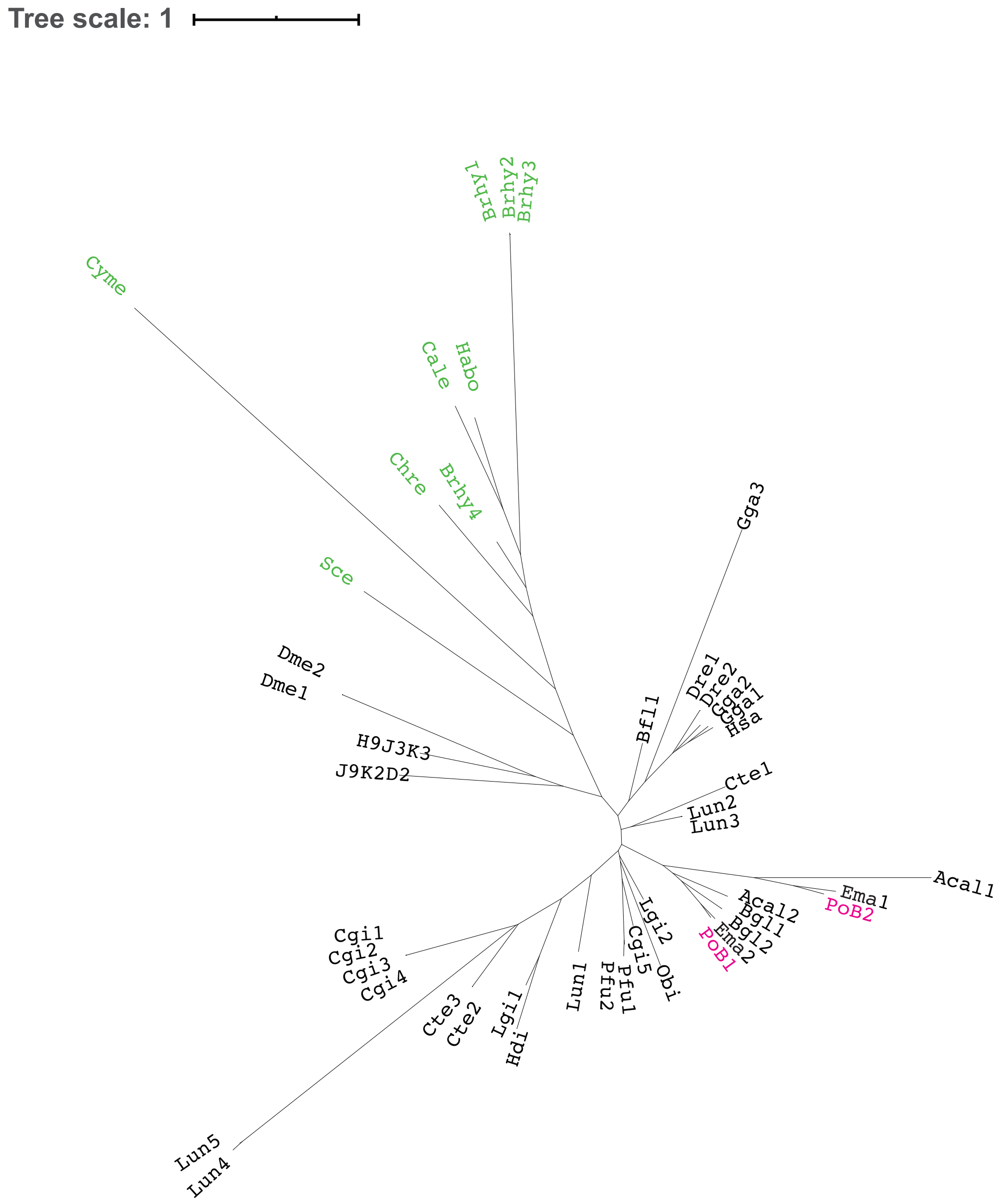


Api1; J9K2D2
Acal1; XP_005099547.2
Acal2; XP_012941160.1
Bfl1; C3YX59
Bgl1; XP_013086066.1
Bgl2; XP_013086068.1
Bmo; H9J3K3
Brhy1; B_m.25540
Brhy2; B_m.25550
Brhy3; B_m.25551
Brhy4; B_m.25561
Cte1; R7TC54
Cte2; R7VEA1
Cte3; R7TRQ8
Cale; g2233.t1
Chre; A8IFY3
Cgi1; XP_011433686.1
Cgi2; XP_019924656.1
Cgi3; XP_019924657.1
Cgi4; XP_019924658.1
Cgi5; XP_019929326.1
Cyme; tr_M1V583_M1V583_CYAM1
Dme1; Q95TG7
Dme2; A8DYB0
Ema1; e4271c37.7
Ema2; e4271c37.1
Habo; H_m.52286
Hdi; HDSC49952CG00010
Lun1; A0A1S3H4V0
Lun2; A0A1S3H6C9
Lun3; A0A1S3IXZ2
Lun4; A0A1S3H7D6
Lun5; A0A1S3H4I4
Lgi1; XP_009045458.1
Lgi2; XP_009045461.1
Obi; XP_014789895.1
PoB1; p258757c71.61
PoB2; p258757c71.57
Sce; YLR106C
Dre1; E7F4D6
Dre2; F1QMM1
Gga1; E1C4X6
Gga2; A0A1D5P4E1
Gga3; A0A1D5PNK8
Hsa; Q9NU22
Pfu1; aug2.0_176.1_20367.t1
Pfu2; aug2.0_10247.1_26360.t1
