## Supplementary Figures for "Chloroplast acquisition without the gene transfer in kleptoplastic sea slugs, *Plakobranchus ocellatus*": Figure_4_supple_7.pdf

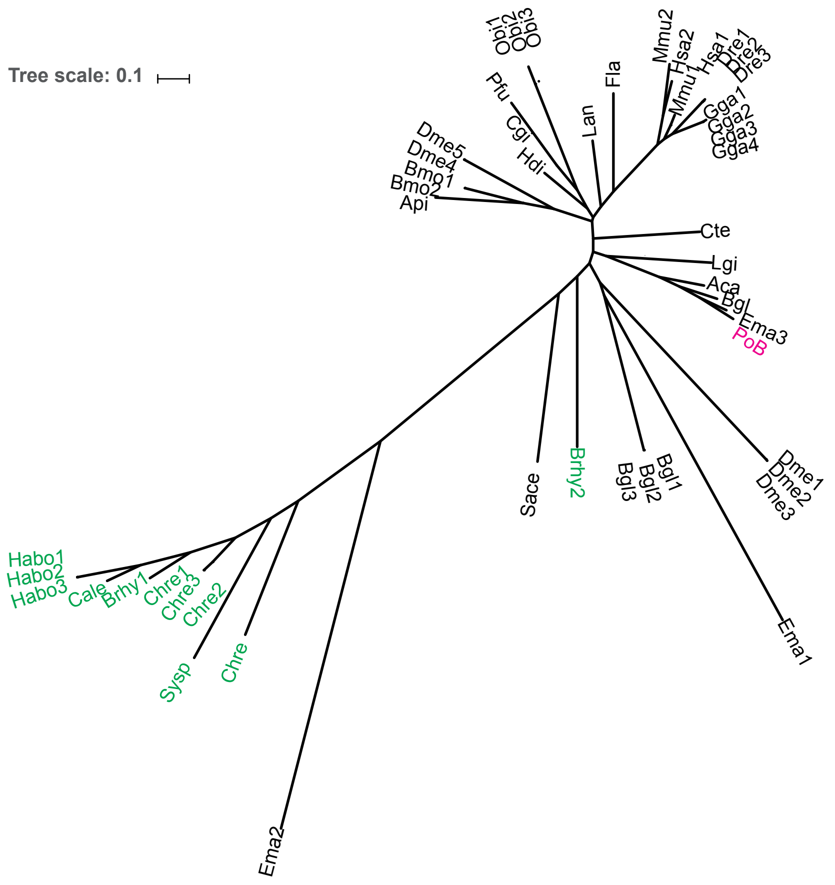


Aca; XP 012936693.1
Api; J9K4W4
Bgl; XP 013066033.1
Bgl1; XP 013061476.1
Bgl2; XP 013061475.1
Bgl3; XP 013061477.1
Bmo1; H9JDT4
Bmo2; E5EVW6
Cgi; XP 011419255.1
Cte; R7TJM0
Dme1; Q8T447
Dme2; B5RJA1
Dme3; Q9VQF4
Dme4; Q01604
Dme5; M9PCE0
Dre1; F1QXV8
Dre3; Q6P003
Dre2; Q7ZV29
Ema1; e11561c44.20
Ema2; e21315c69.33
Ema3; e3214c43.1
Fla; C3XW30
Gga1; P51903
Gga2; A0A1L1RXH1
Gga3; A0A1D5NZW9
Gga4; F1NU17
Hdi; HDSC00324CG00210
Hsa1; P00558
Hsa2; P07205
Lan; A0A1S3HH77
Lgi; XP 009047142.1
Mmu1; P09411
Mmu2; P09041
Obi1; XP 014774198.1
Obi2; XP 014774182.1
Obi3; XP 014774190.1
Pfu; aug2.0 329.1 30622.t1
PoB; p105c62.89

Brhy1; B_m.48831
Brhy2; B_m.33737
Chre; tr A0A125YSW1
Chre1; Q548U3
Chre2; P41758
Chre3; A8JC04
Cale; g2055.t1
Habo1; H_m.33221
Habo2; H_m.33202
Habo3; H_m.33212
Sace; YCR012W
Sysp; P74421 PGK SYNY3
