## Supplementary Figures for "Chloroplast acquisition without the gene transfer in kleptoplastic sea slugs, *Plakobranchus ocellatus*": Figure_4_supple_8.pdf

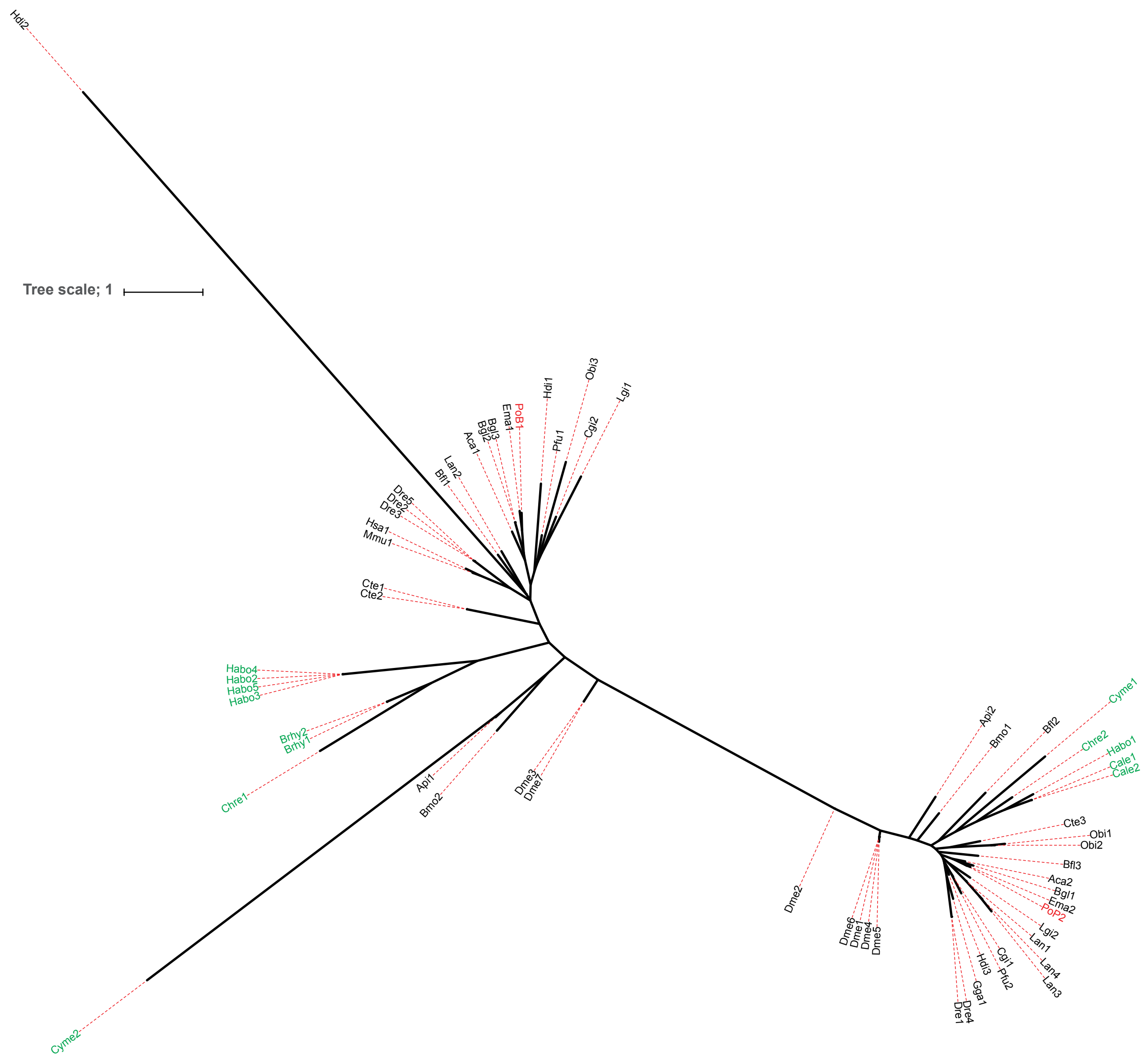


Dre1; A0A0R4IS22
Lan1; A0A1S3H867
Lan2; A0A1S3HV36
Lan3; A0A1S3K7R9
Lan4; A0A2R2MN91
Dme1; A1Z757
Dme2; A1Z758
Dme3; A1Z759
Chre1; A8JAN4
Pfu1; aug2.0_4854.1_16038.t1
Pfu2; aug2.0_680.1_01048.t1
Bfl1; C3Y1S2
Bfl2; C3ZD92
Bfl3; C3ZWY3
Ema1; e2072c39.23
Ema2; e9727c36.9
PoB1; p440c72.239
PoB2; p855c67.9
Gga1; F1NSZ2
Dre2; F1Q8W7
Bmo1; H9J281
Bmo2; H9JWI3
Hdi1; HDSC00027CG00110
Hdi2; HDSC04373CG00010
Hdi3; HDSC07471CG00010
Api1; J9JT18
Api2; J9K3D6

Dme4; Q24443
Dme5; Q26373
Dre3; Q3B7R6
Mmu1; Q562D6
Dme6; Q7JY97
Dre4; Q7SYI9
Dme7; Q8MR50
Hsa1; Q9BU70
Chre2; Q9XHE2
Dre5; R4GEK9
Cte1; R7T6M9
Cte2; R7THN0
Cte3; R7TV65
Aca1; XP_005089019.1
Lgi1; XP_009048710.1
Lgi2; XP_009060728.1
Cgi1; XP_011422754.2
Cgi2; XP_011442441.2
Aca2; XP_012935158.1
Bgl1; XP_013068586.1
Bgl2; XP_013073365.1
Bgl3; XP_013073366.1
Obi1; XP_014780460.1
Obi2; XP_014780461.1
Obi3; XP_014781937.1

Habo1; H_m.22572
Habo2; H_m.23039
Habo3; H_m.23046
Habo4; H_m.23050
Habo5; H_m.23056
Cale1; g7280.t1
Cale2; g7280.t2
Cyme1; tr_M1V506_CYAM1
Cyme2; tr_M1VLH0_CYAM1
Brhy1; B_m.26666
Brhy2; B_m.26667
