## Supplementary Figures for "Chloroplast acquisition without the gene transfer in kleptoplastic sea slugs, *Plakobranchus ocellatus*": Figure_6_supple_5.pdf

### OG0000005; Lectin group

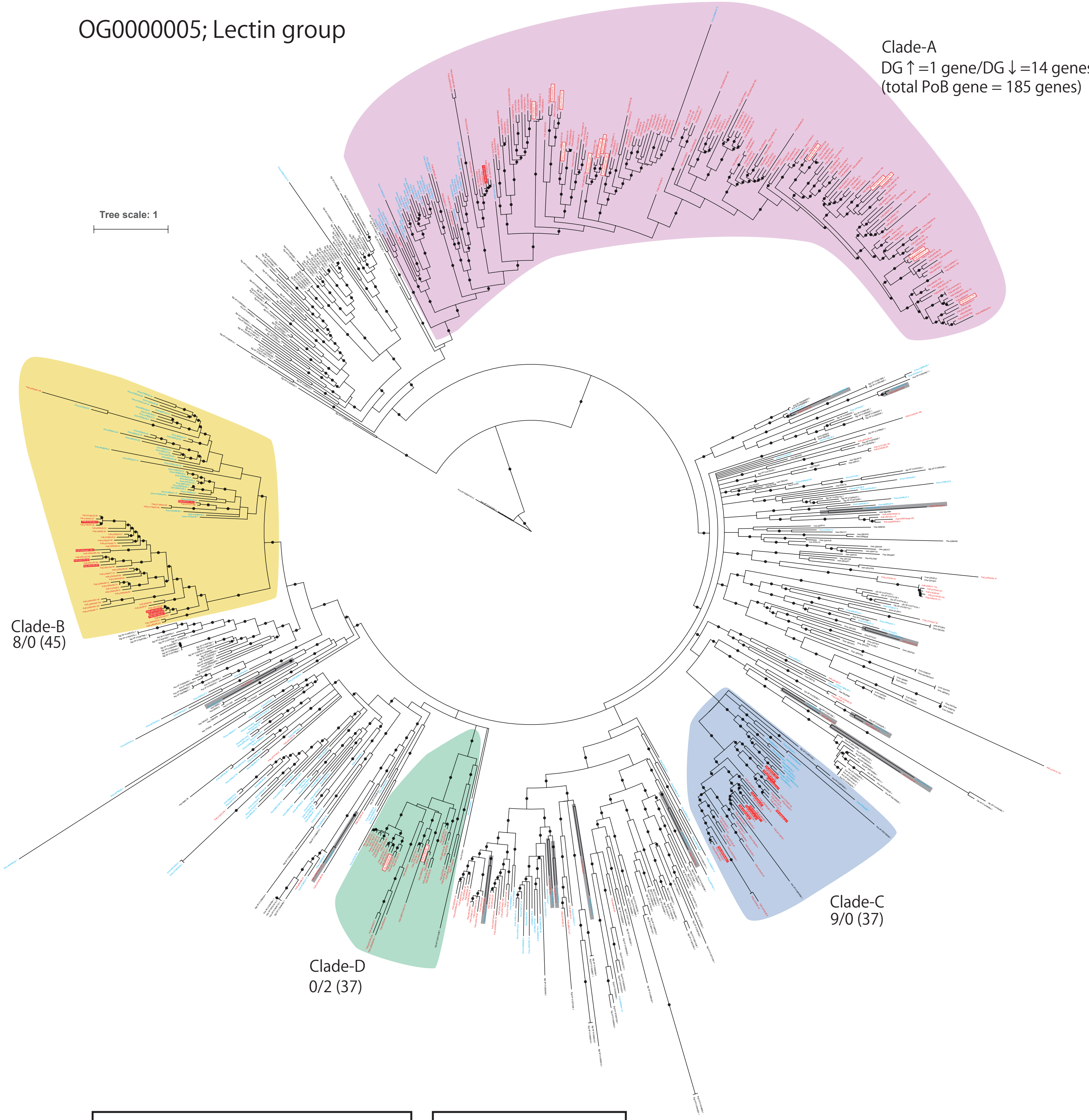


| DG-downregulated genes | DG-upregulated genes |
|---|---|
| p339c68.52 | p3334c67.98 |
| p1147c61.57 | p3054c66.152 |
| p335c60.39 | p501c63.67 |
| p1352c59.54 | p501c63.64 |
| p6313c71.6 | p501c63.56 |
| p124c71.259 | p4334c64.28 |
| p227c62.170 | p47c72.174 |
| p339c68.166 | p501c63.63 |
| p1857c64.5 | p4334c64.26 |
| p1209c50.1 | p501c63.50 |
| p258753c65.149 | p7187c46.2 |
| p1121c63.100 | p4334c64.30 |
| p124c71.261 | p855c67.129 |
| p124c71.266 | p335c60.28 |
| p256c68.16 | p258754c68.4 |
| p1529c64.11 | p258746c66.63 |
| | p958c60.27 |
| | p335c60.25 |

OTU coloration
- PoB
- Ema
- Other species
- DG-upregulated PoB gene
- DG-downregulated PoB gene
