## Supplementary figures and images for "Chloroplast acquisition without the gene transfer in kleptoplastic sea slugs, *Plakobranchus ocellatus*"

### Figure_2_supple_1.jpg

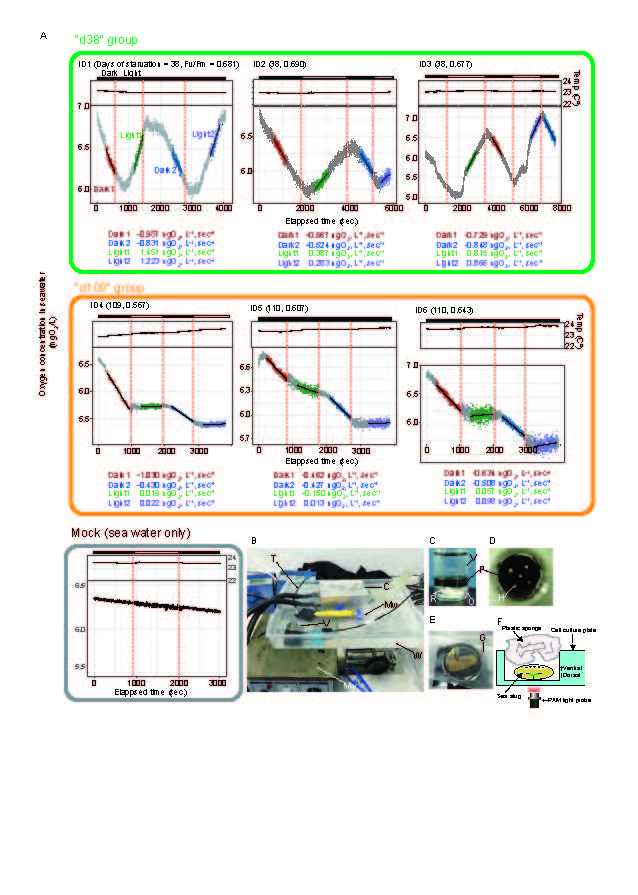

### Figure_3_supple_1.pdf

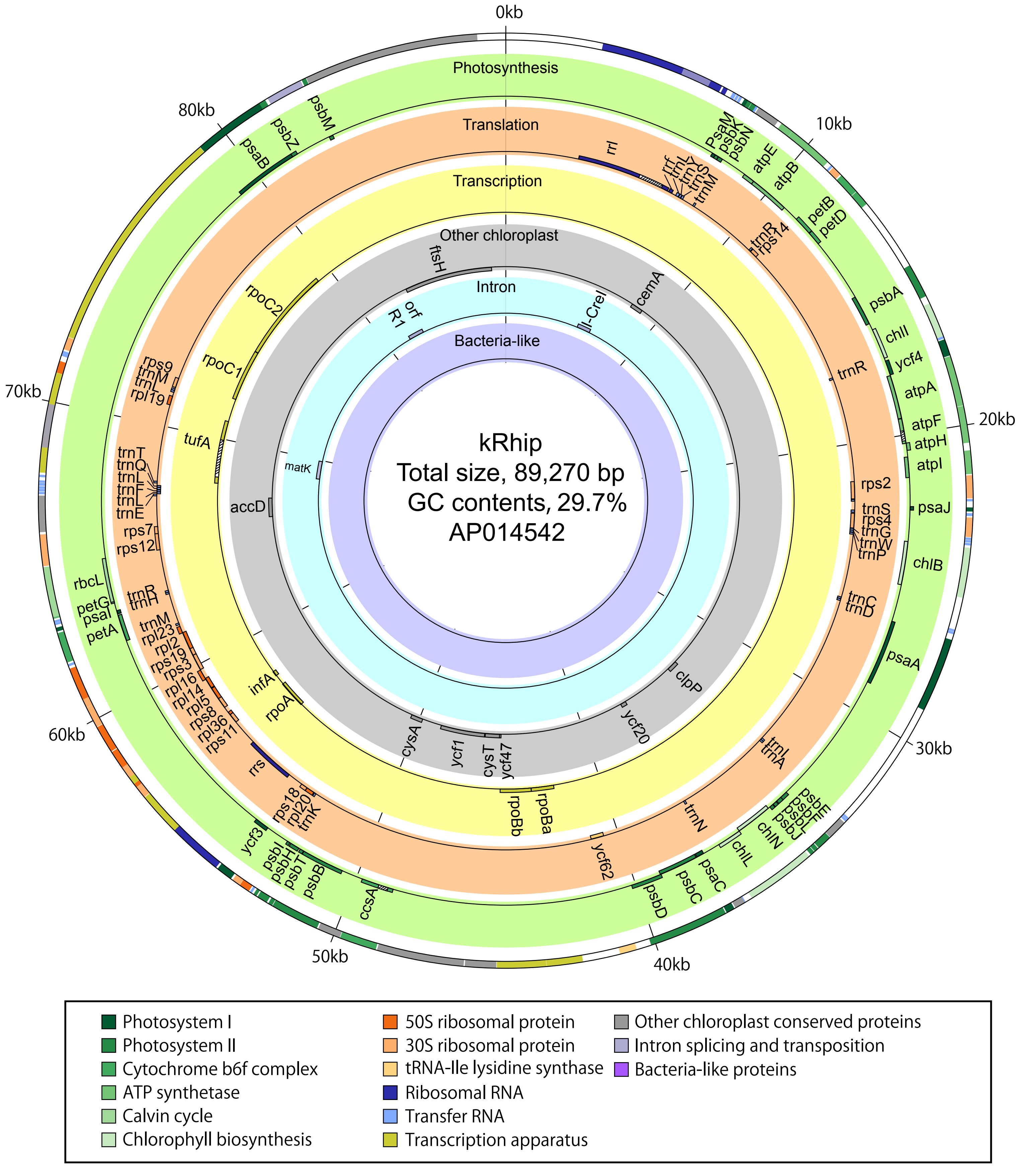

### Figure_3_supple_2.pdf

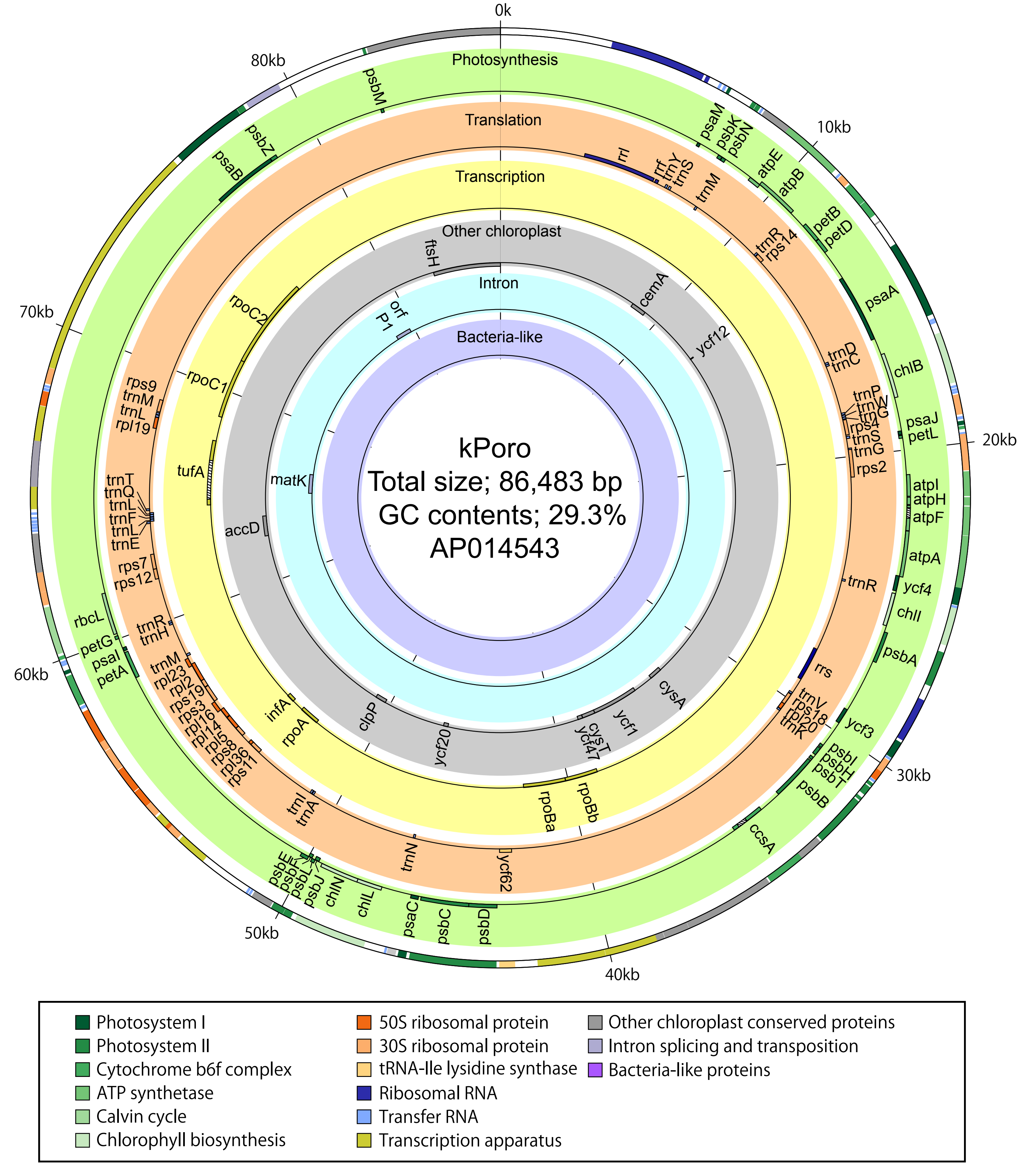

### Figure_3_supple_3.pdf

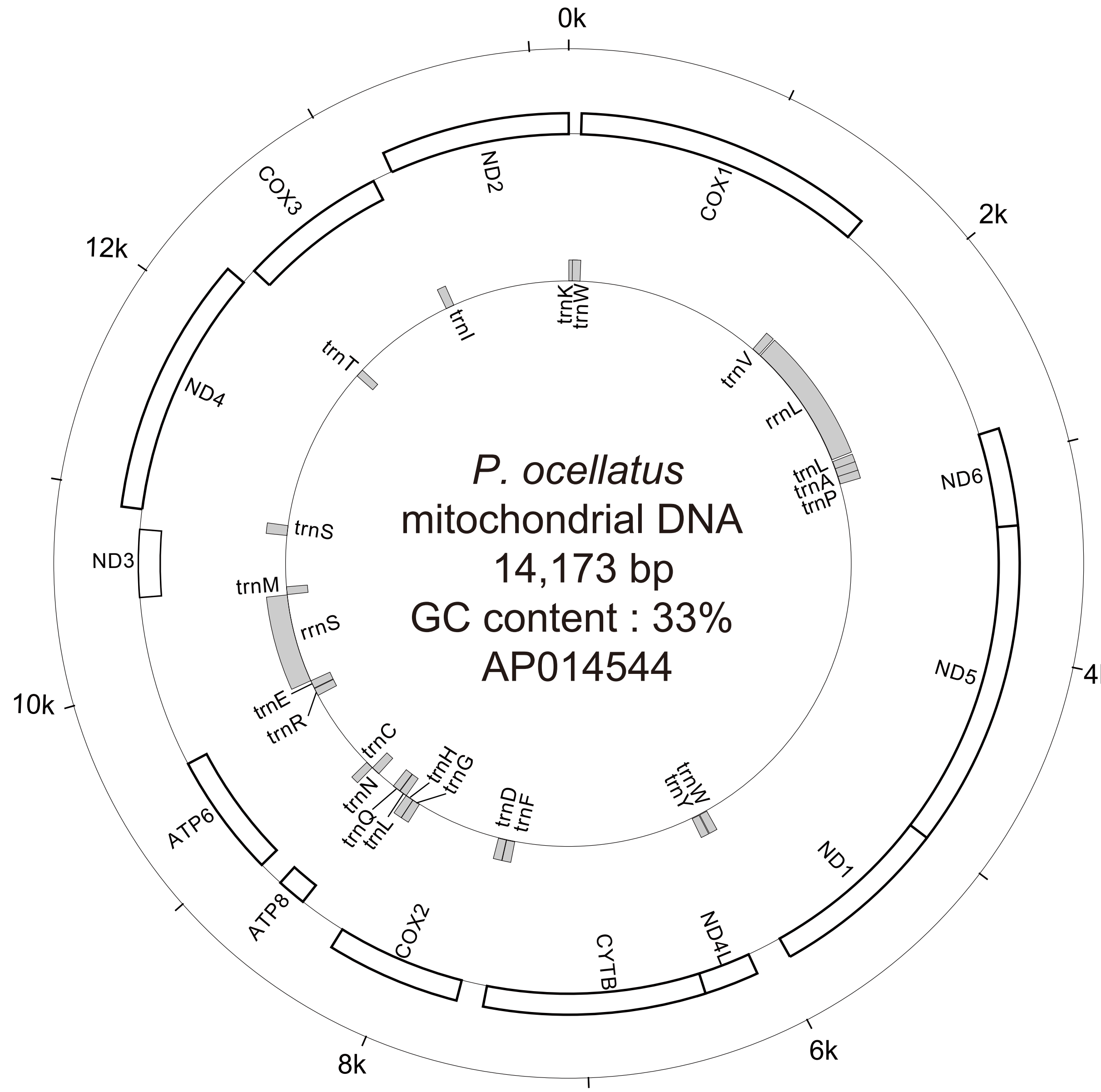

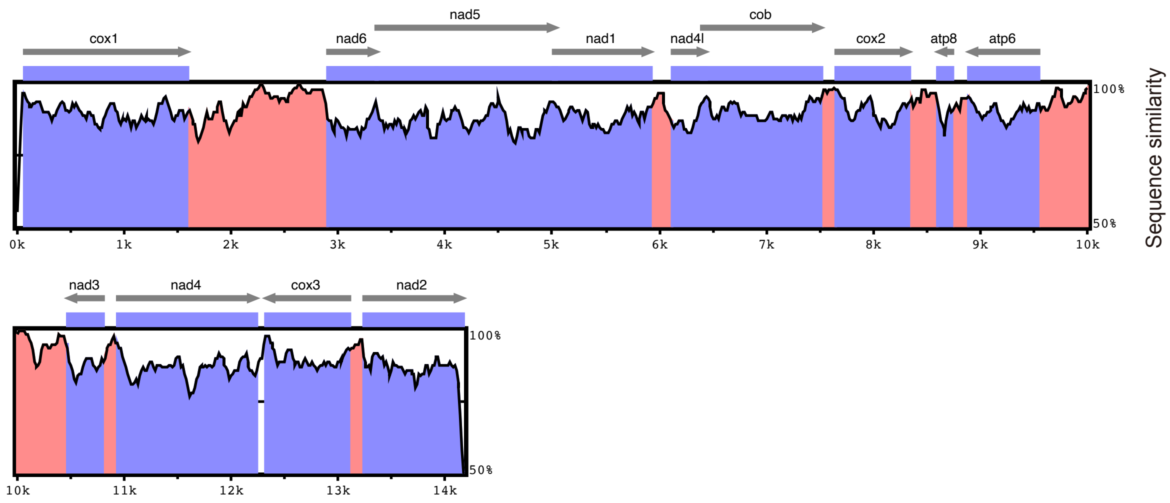

### Figure_3_supple_4.pdf

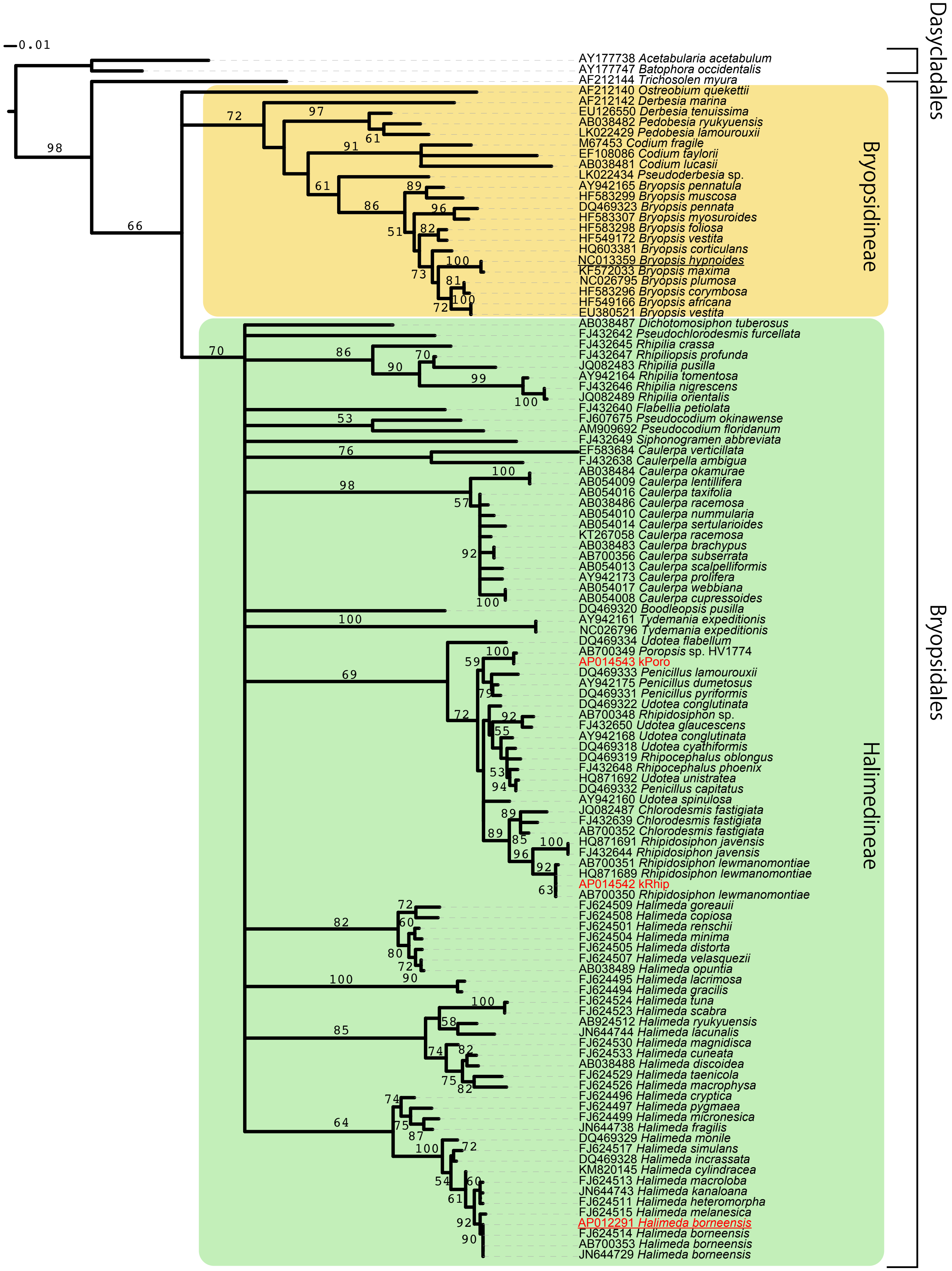

### Figure_3_supple_5.pdf

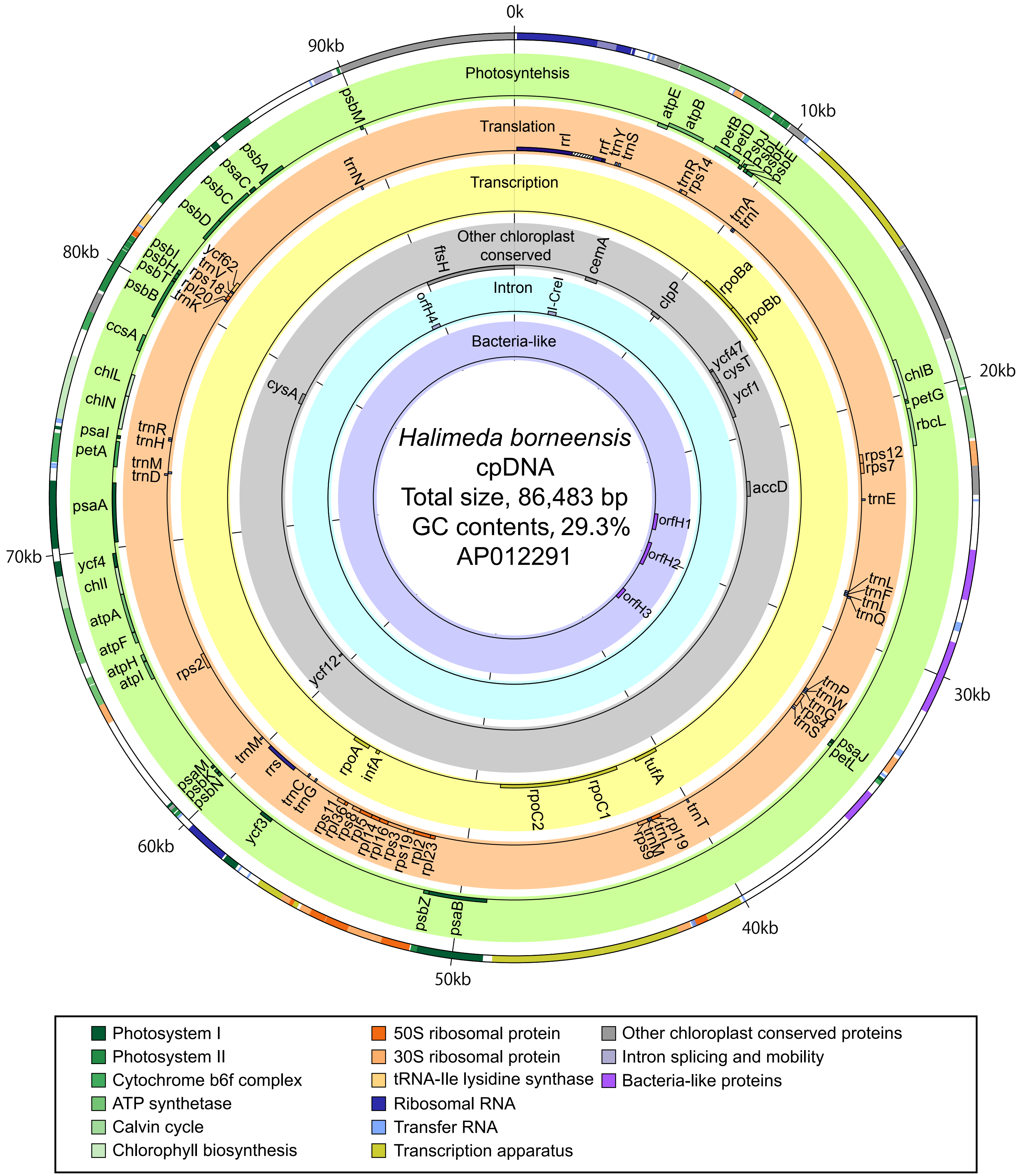

### Figure_3_supple_6.pdf

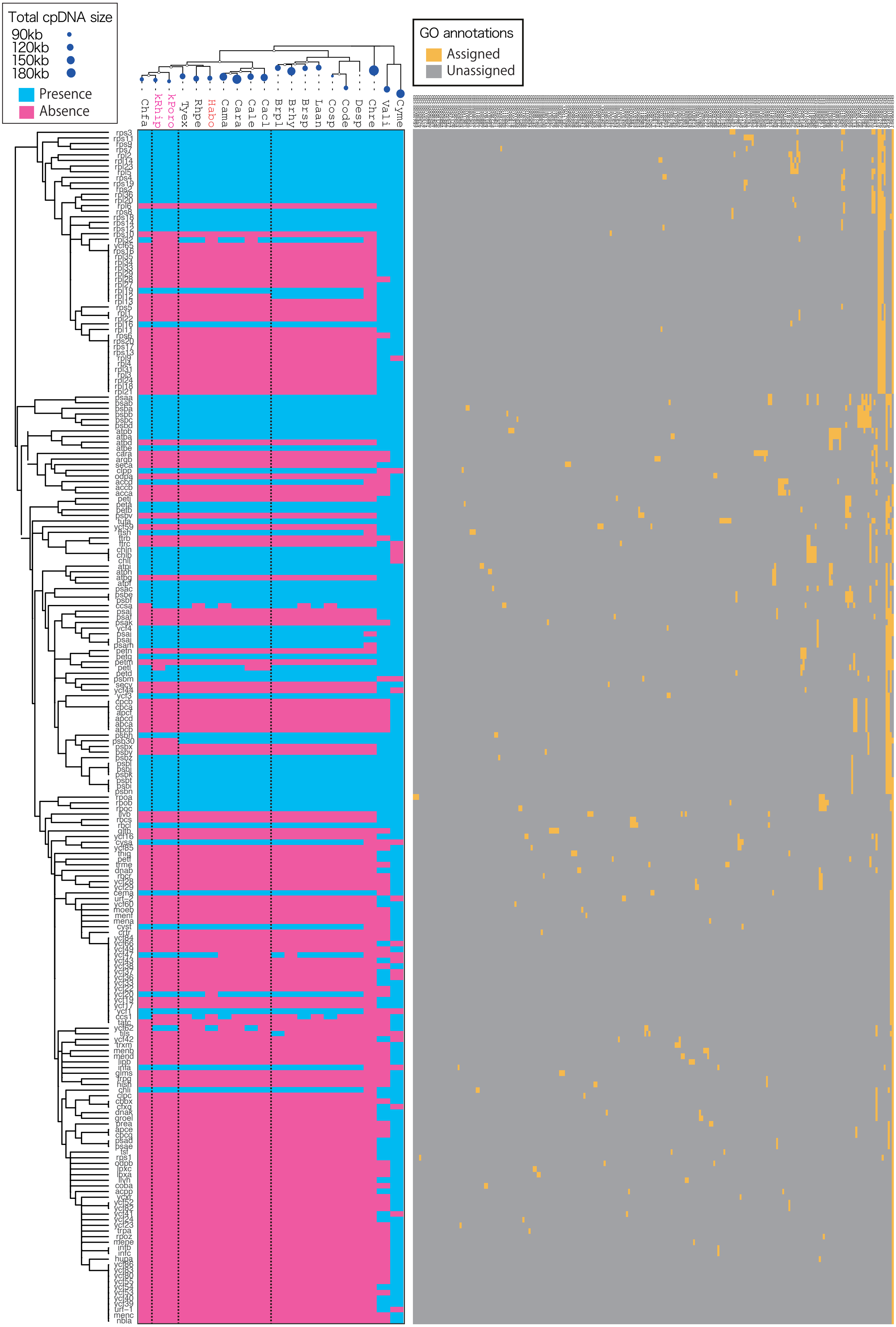

### Figure_3_supple_7.pdf

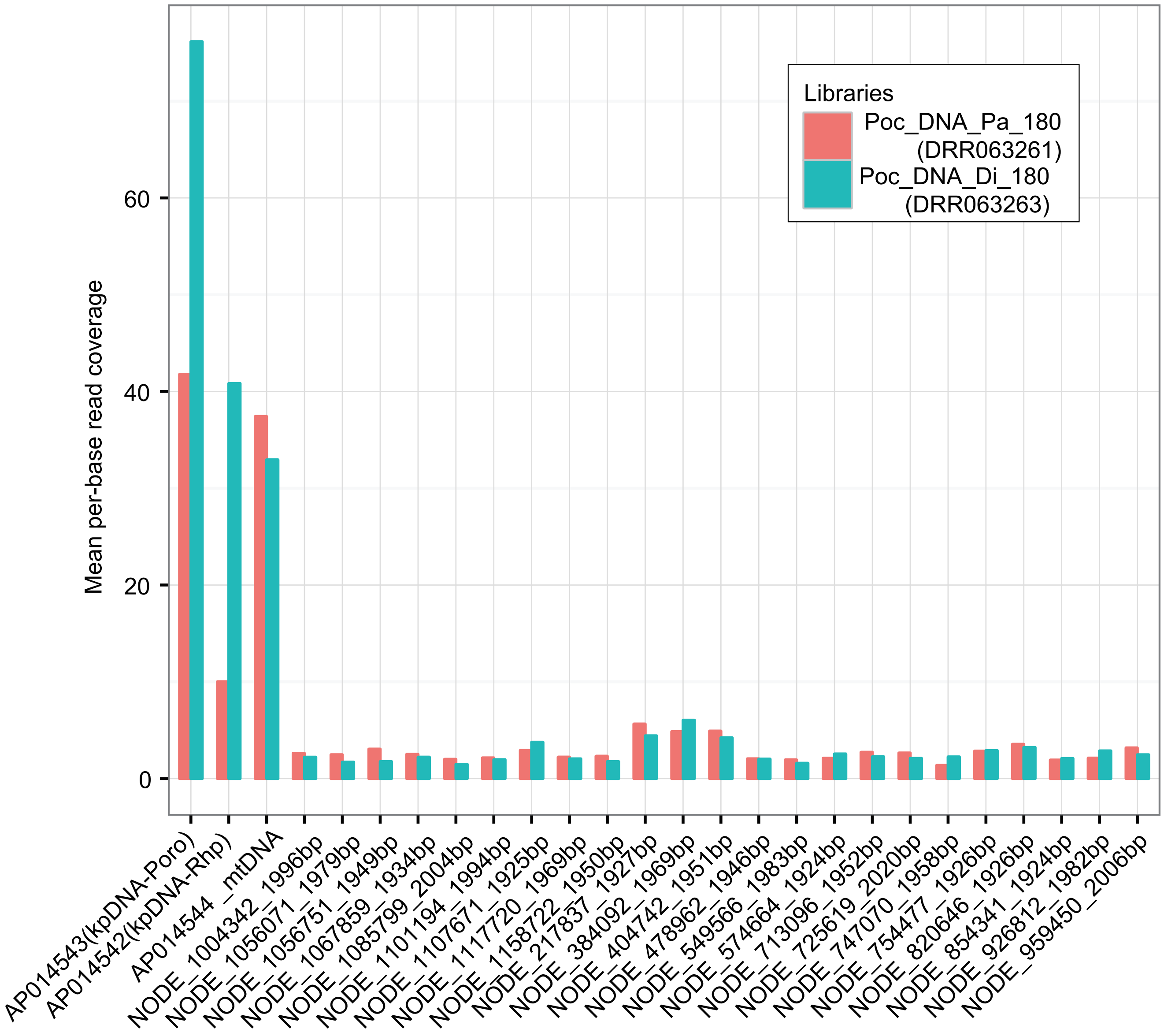

### Figure_3_supple_8.pdf

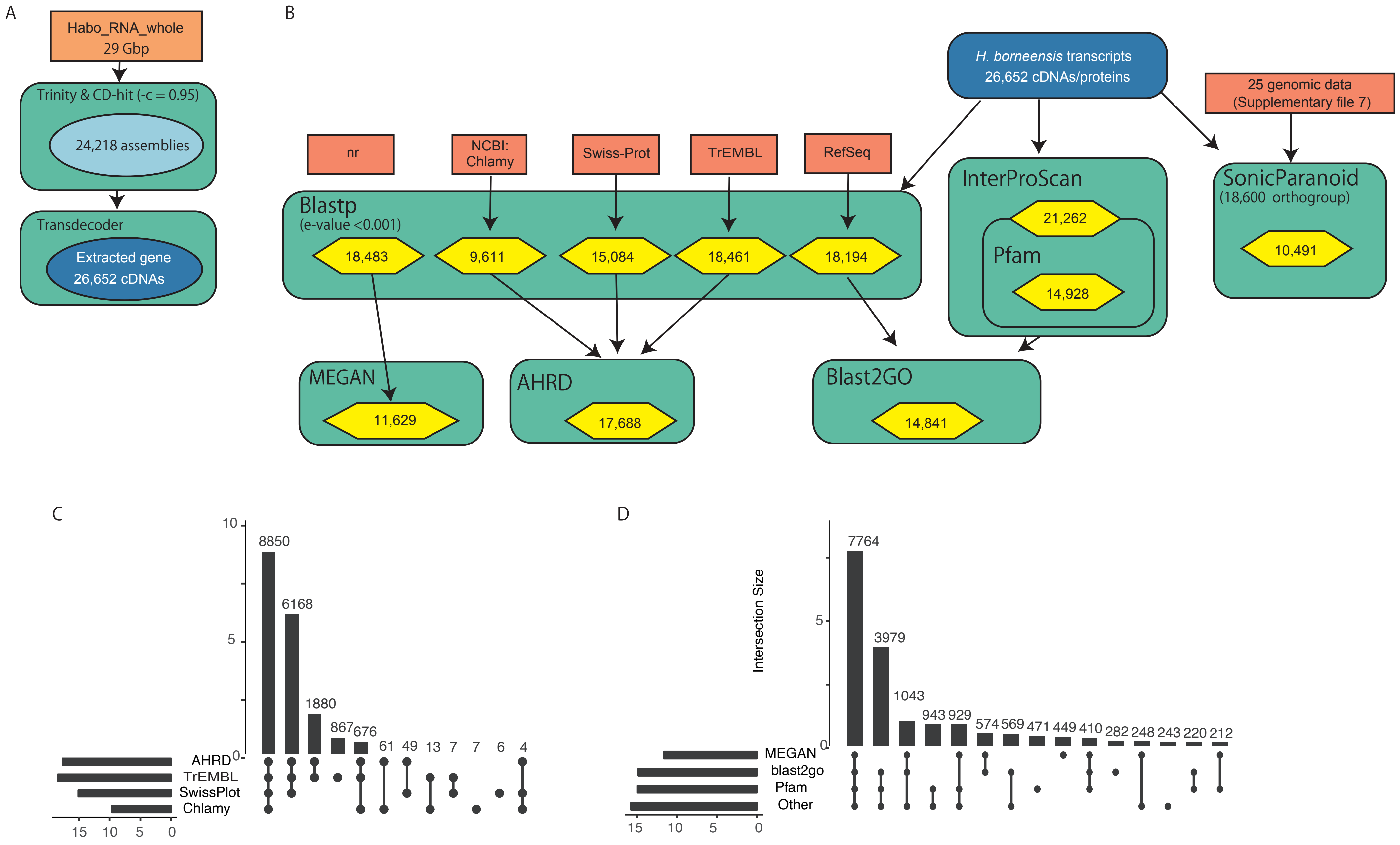

### Figure_3_supple_9.pdf

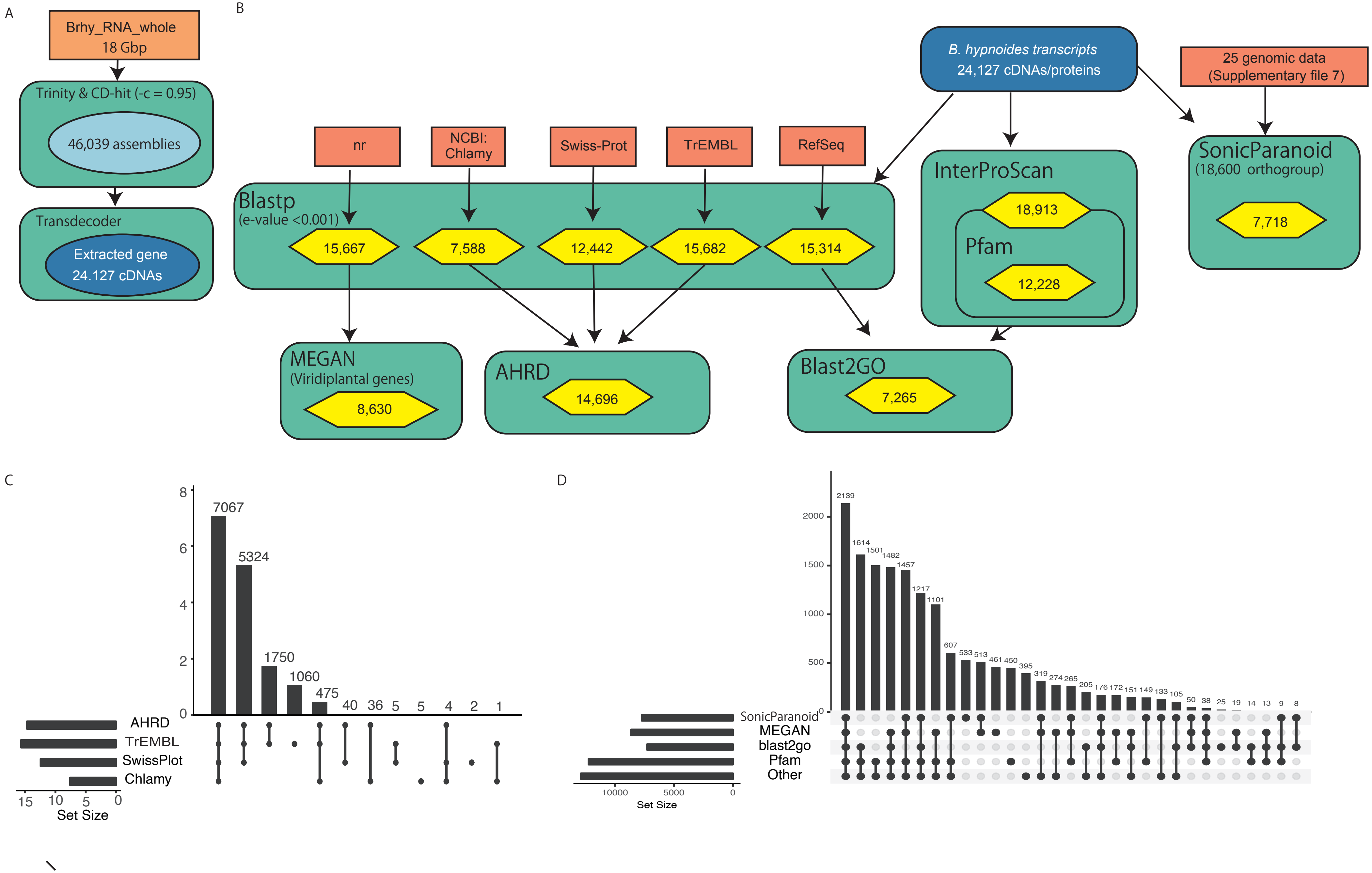

### Figure_3_supple_10.pdf

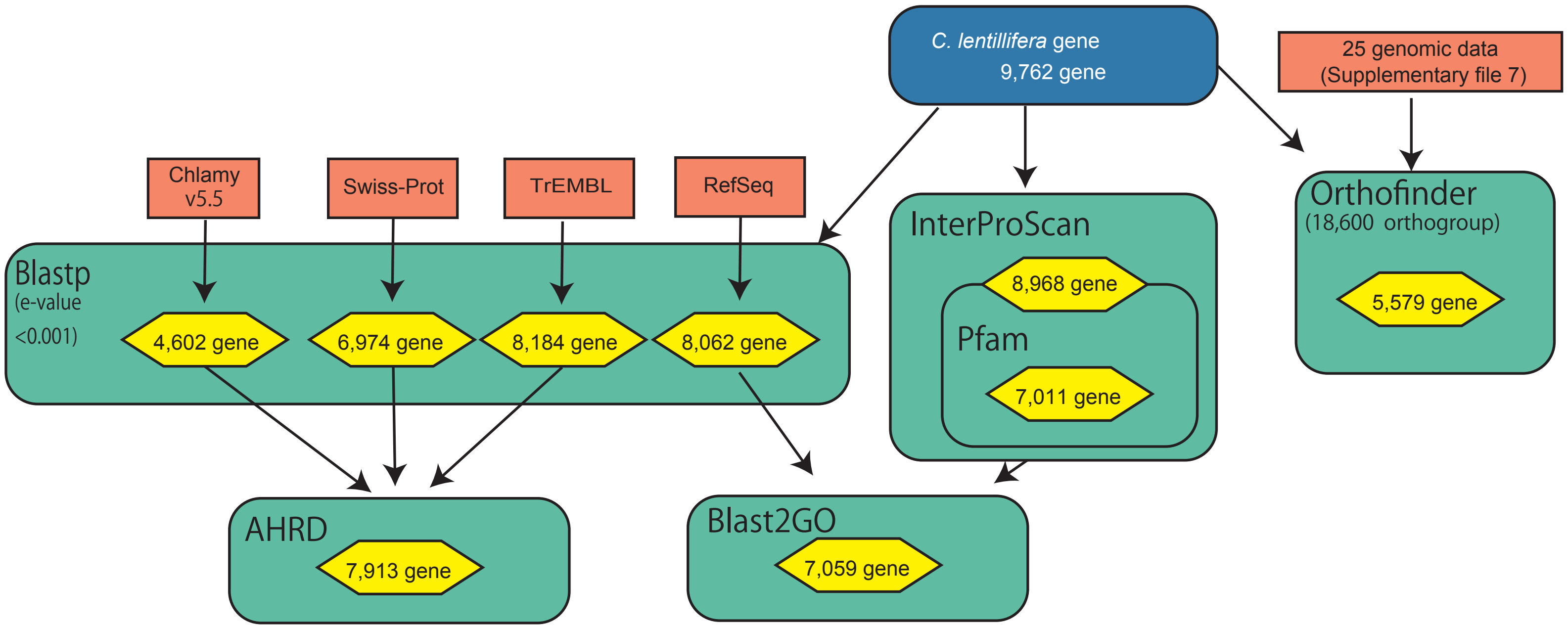

### Figure_4_supple_1.pdf

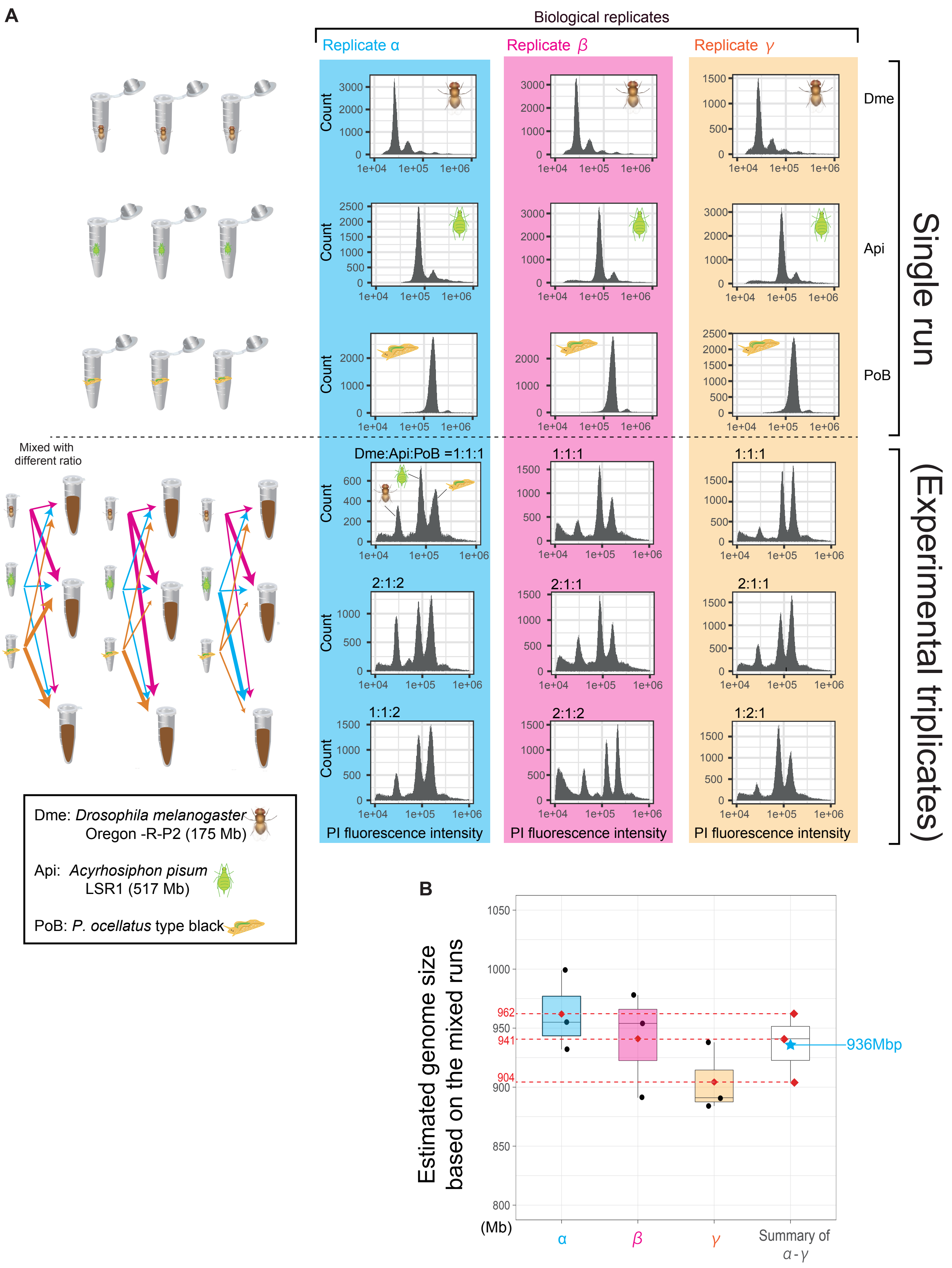

### Figure_4_supple_2.pdf

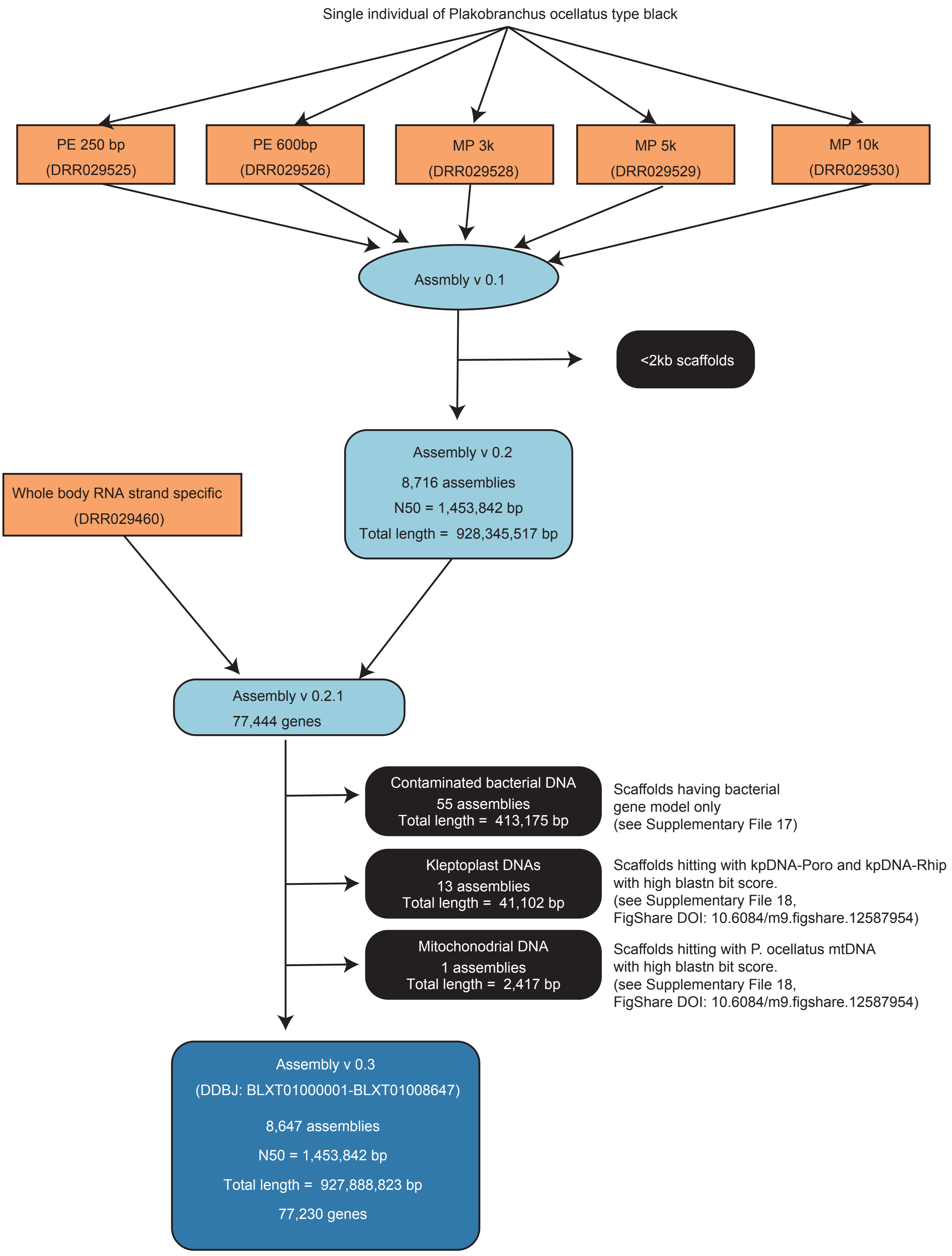

### Figure_4_supple_3.pdf

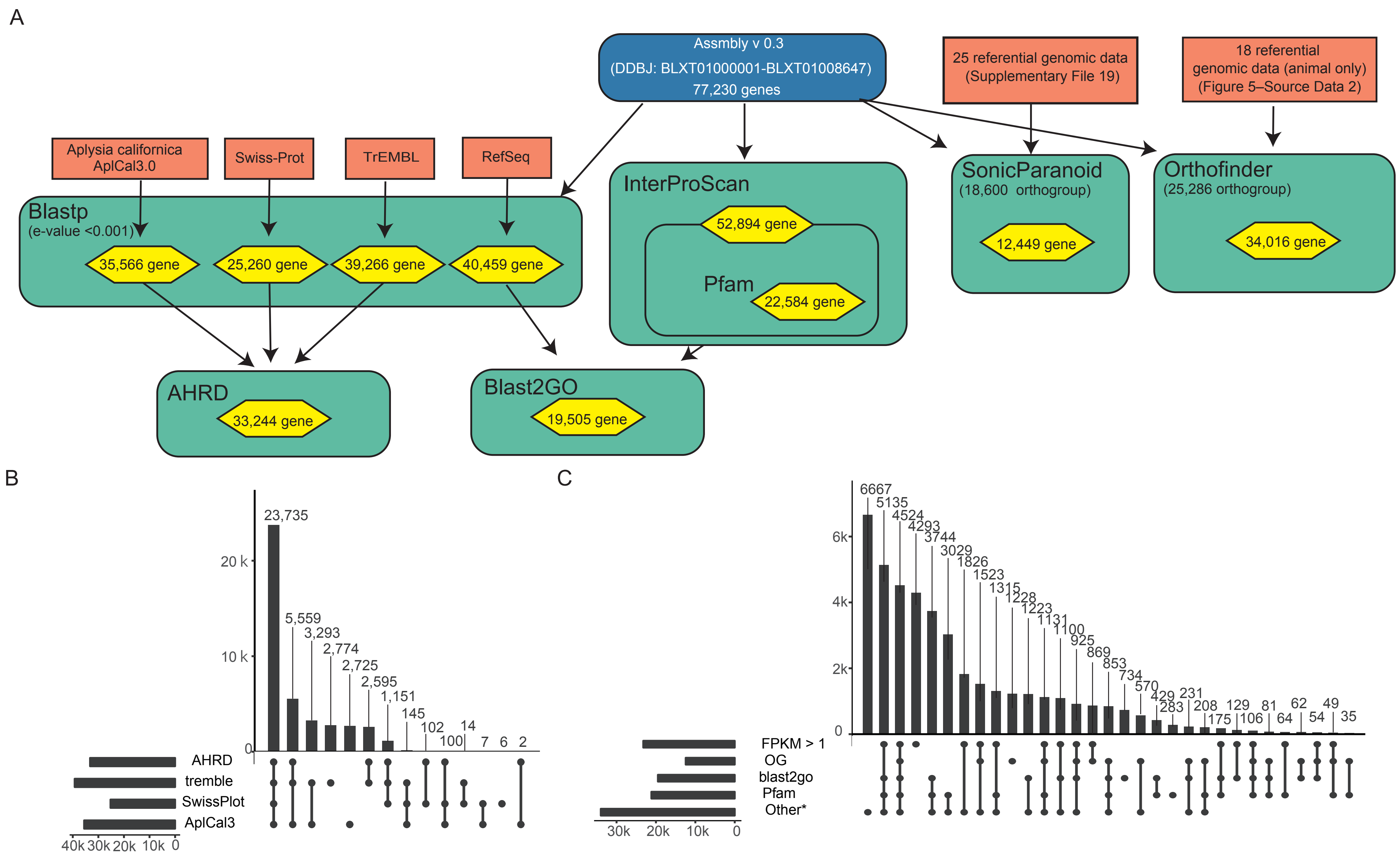

### Figure_4_supple_4.pdf

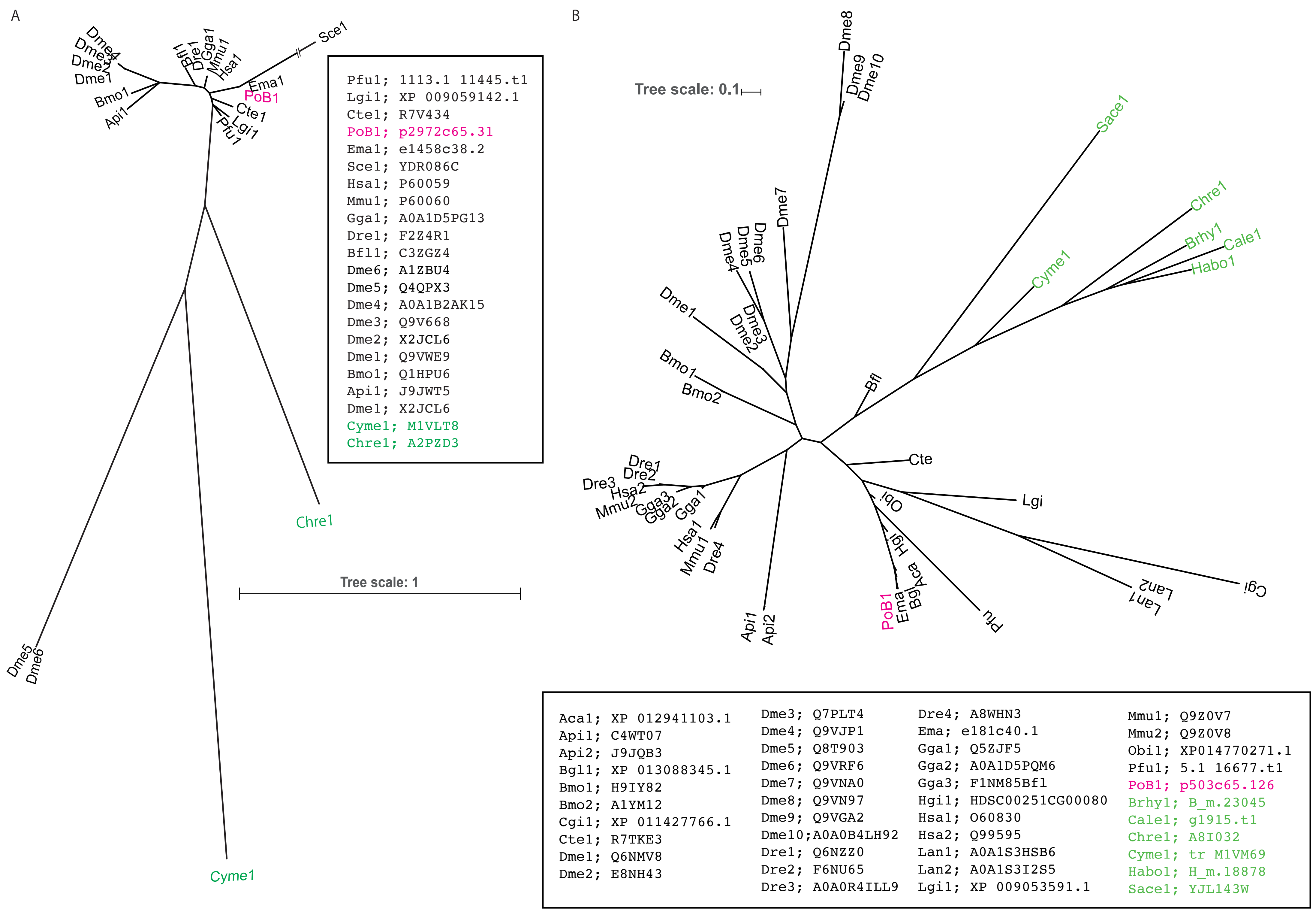

### Figure_4_supple_5.pdf

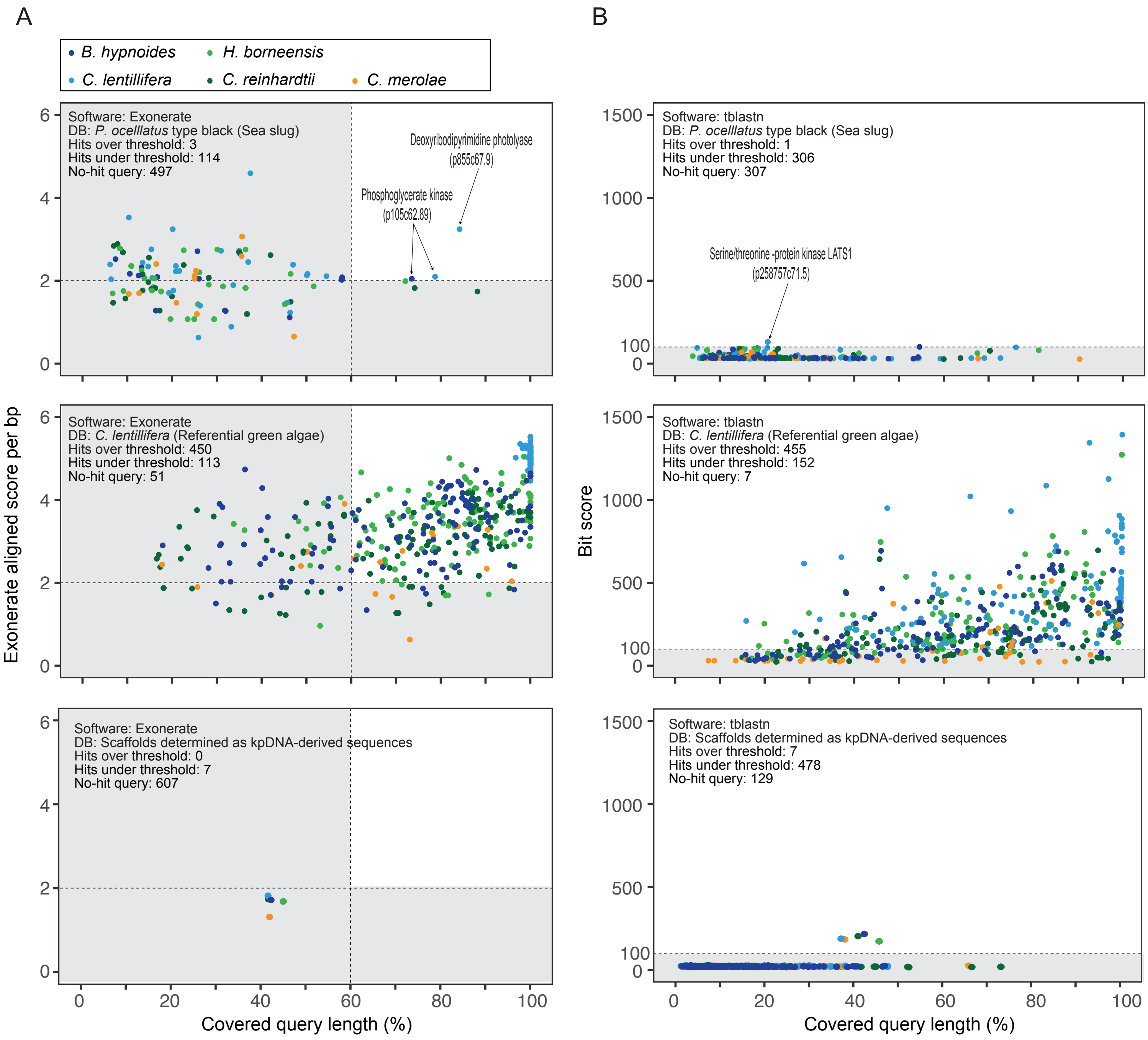

### Figure_4_supple_9.pdf

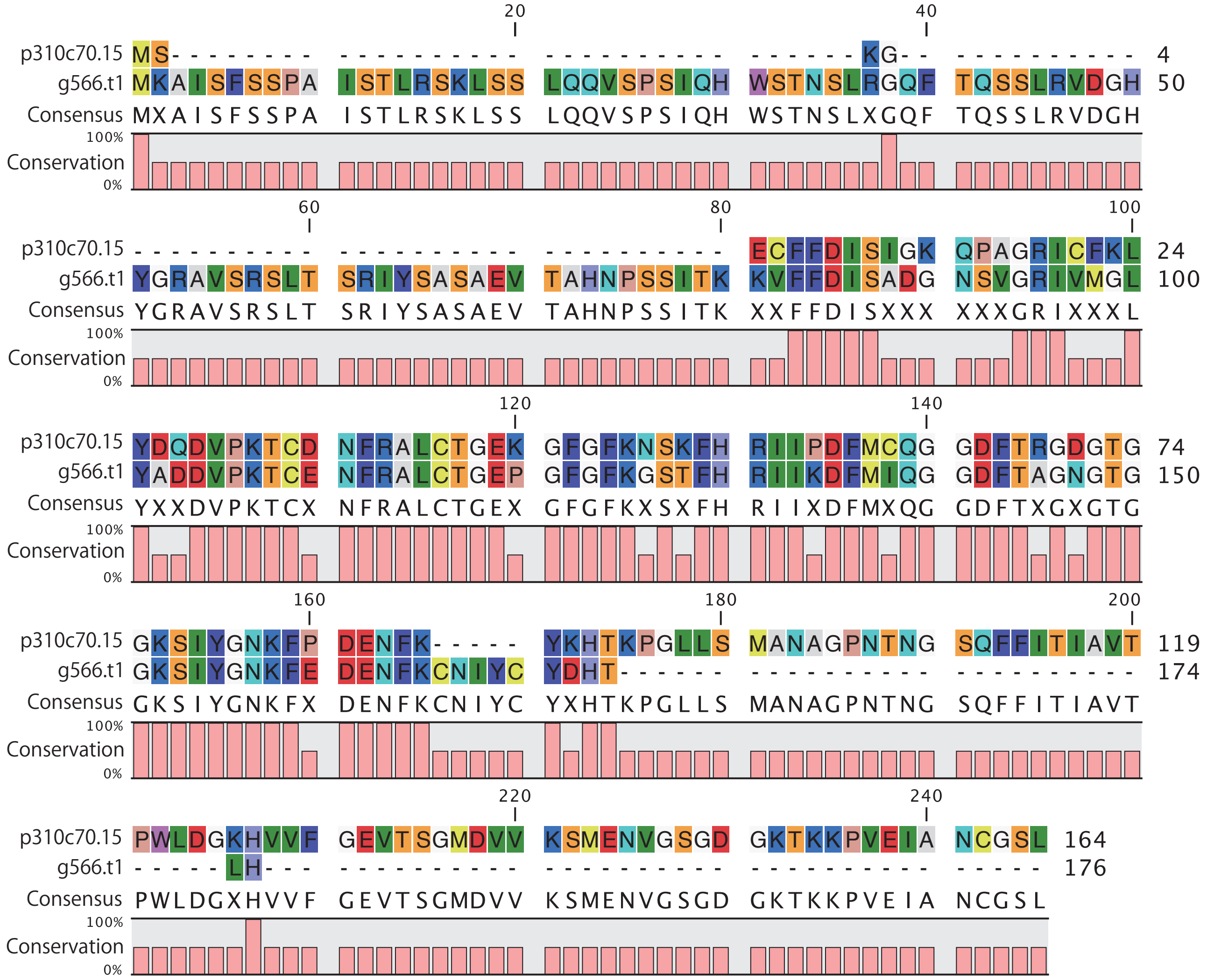

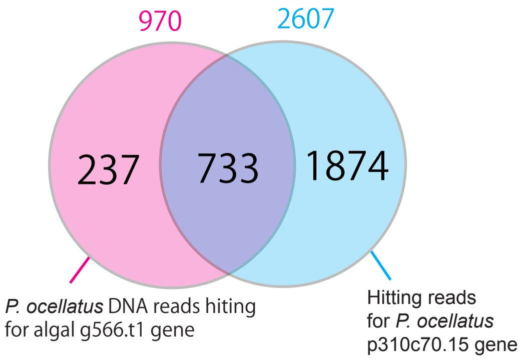

### Figure_4_supple_10.jpg

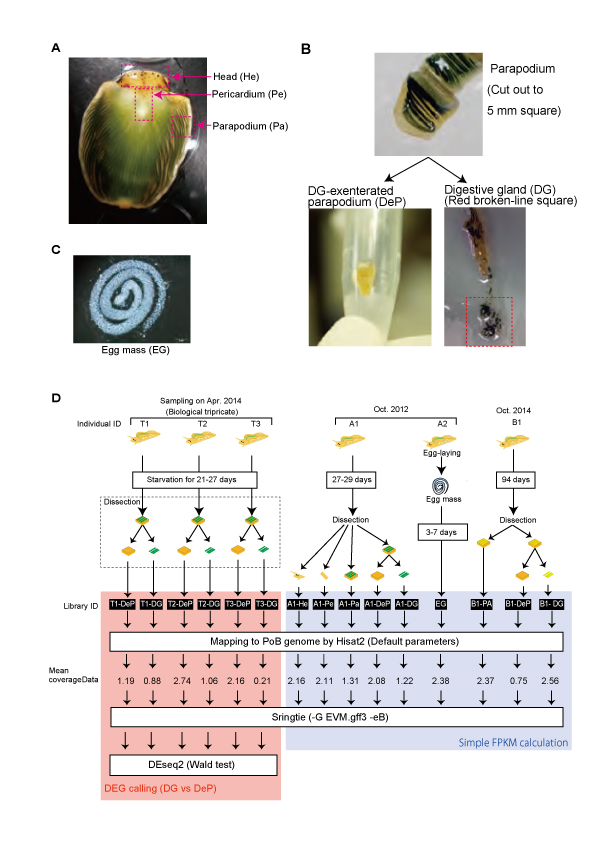

### Figure_4_supple_11.pdf

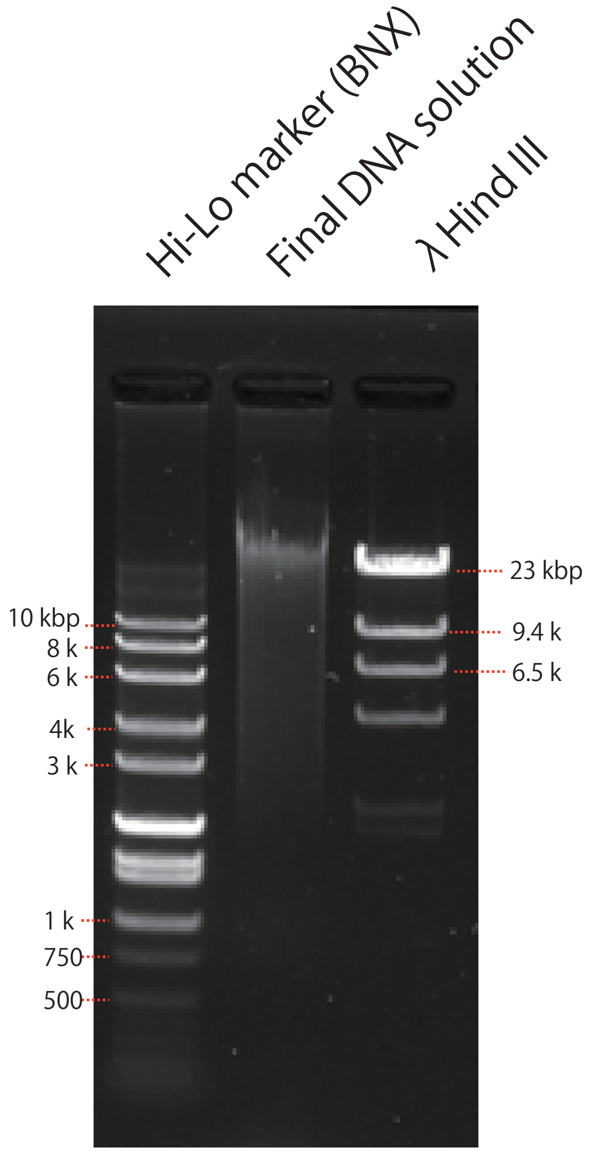

### Figure_4_supple_12.pdf

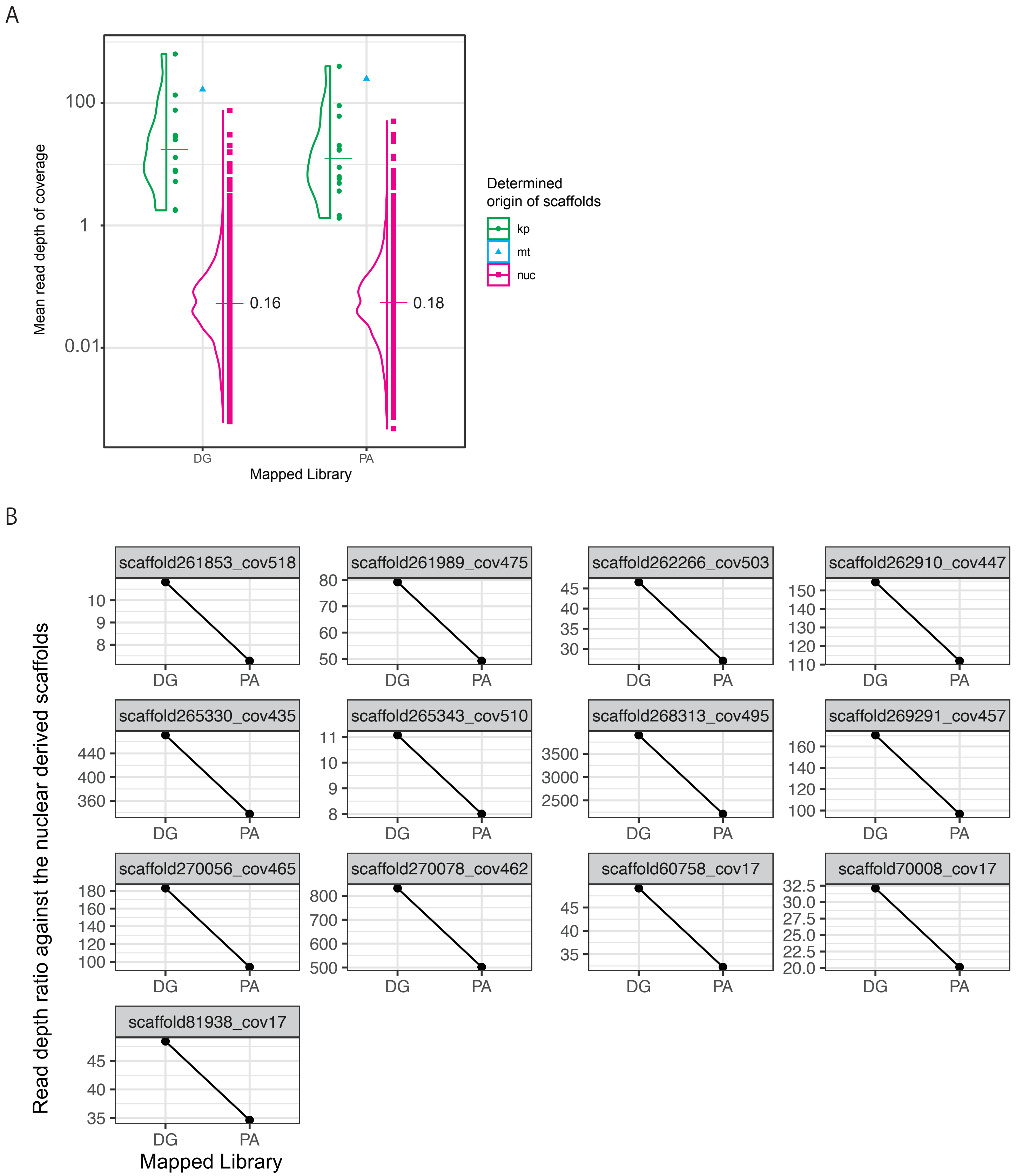

### Figure_4_supple_13.pdf

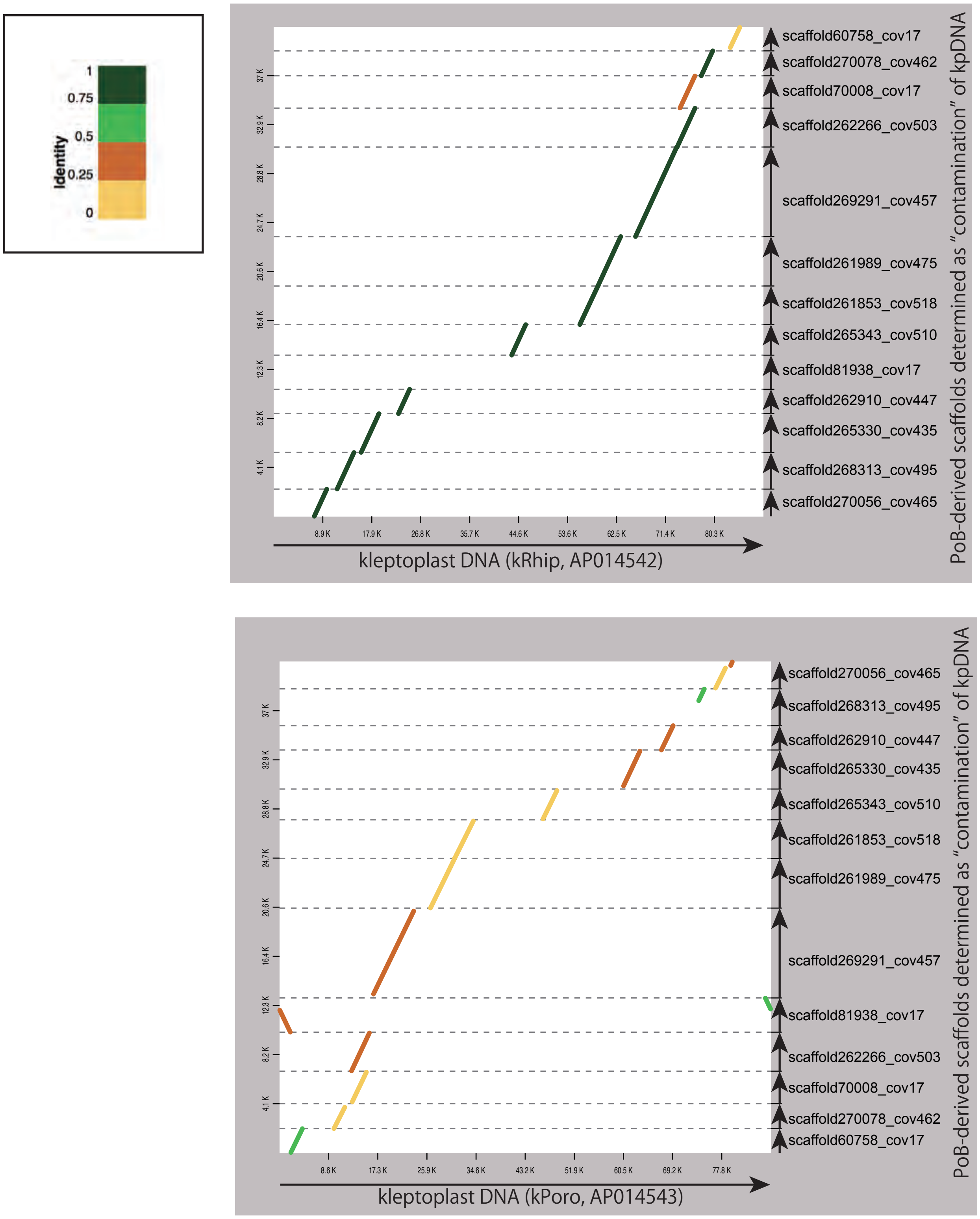

### Figure_4_supple_14.pdf

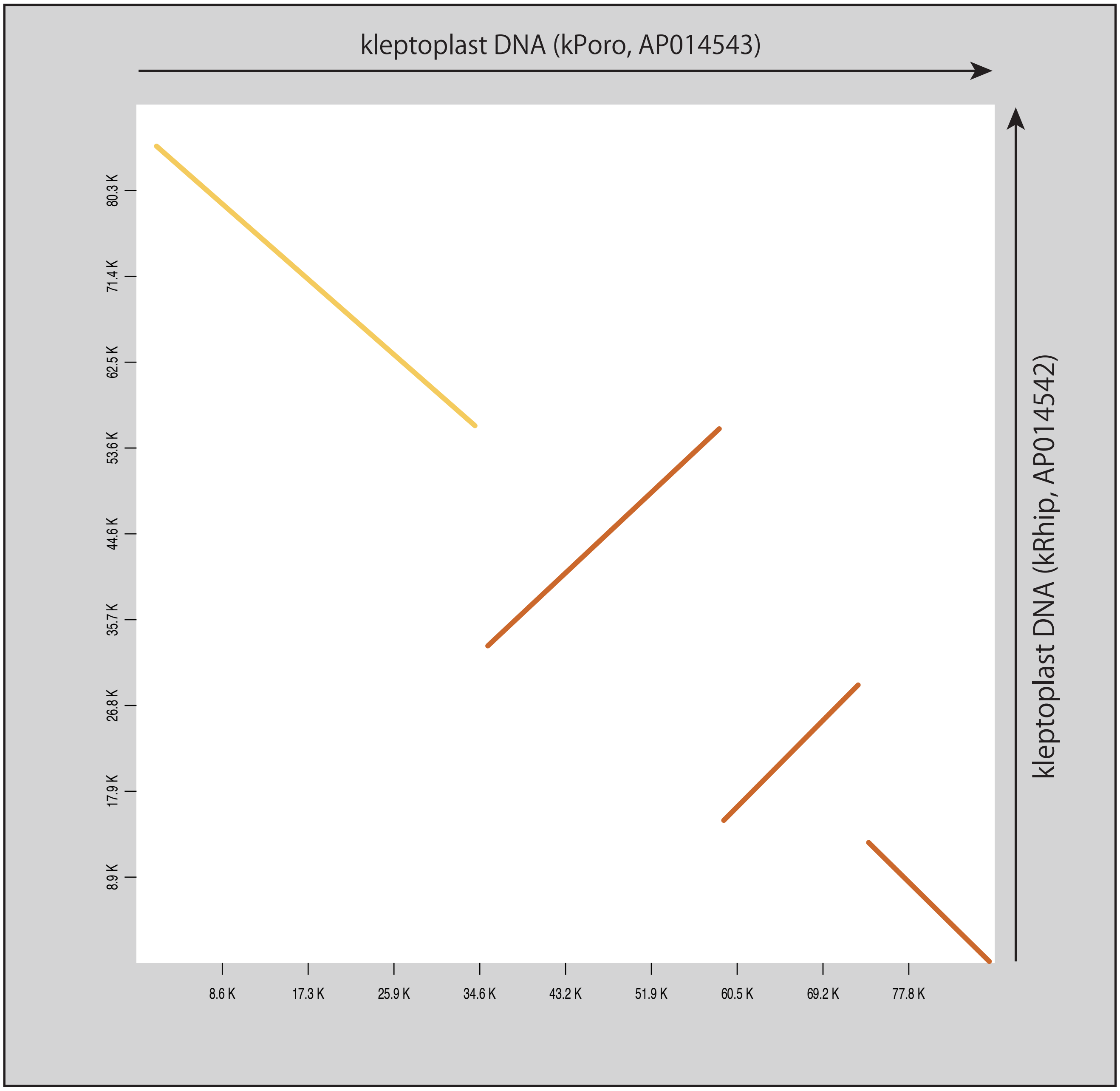

### Figure_4_supple_15.pdf

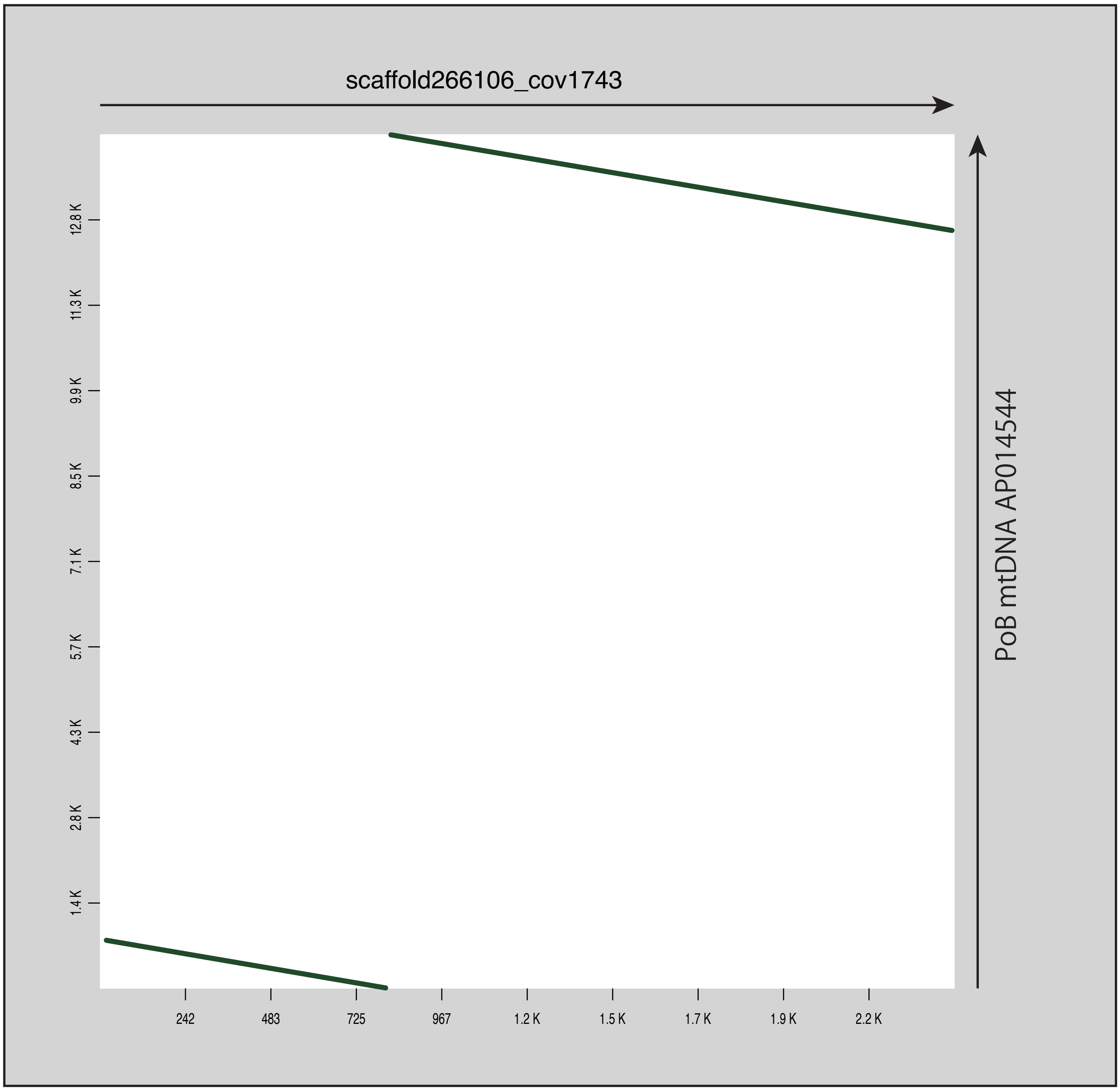

### Figure_5_supple_1.pdf

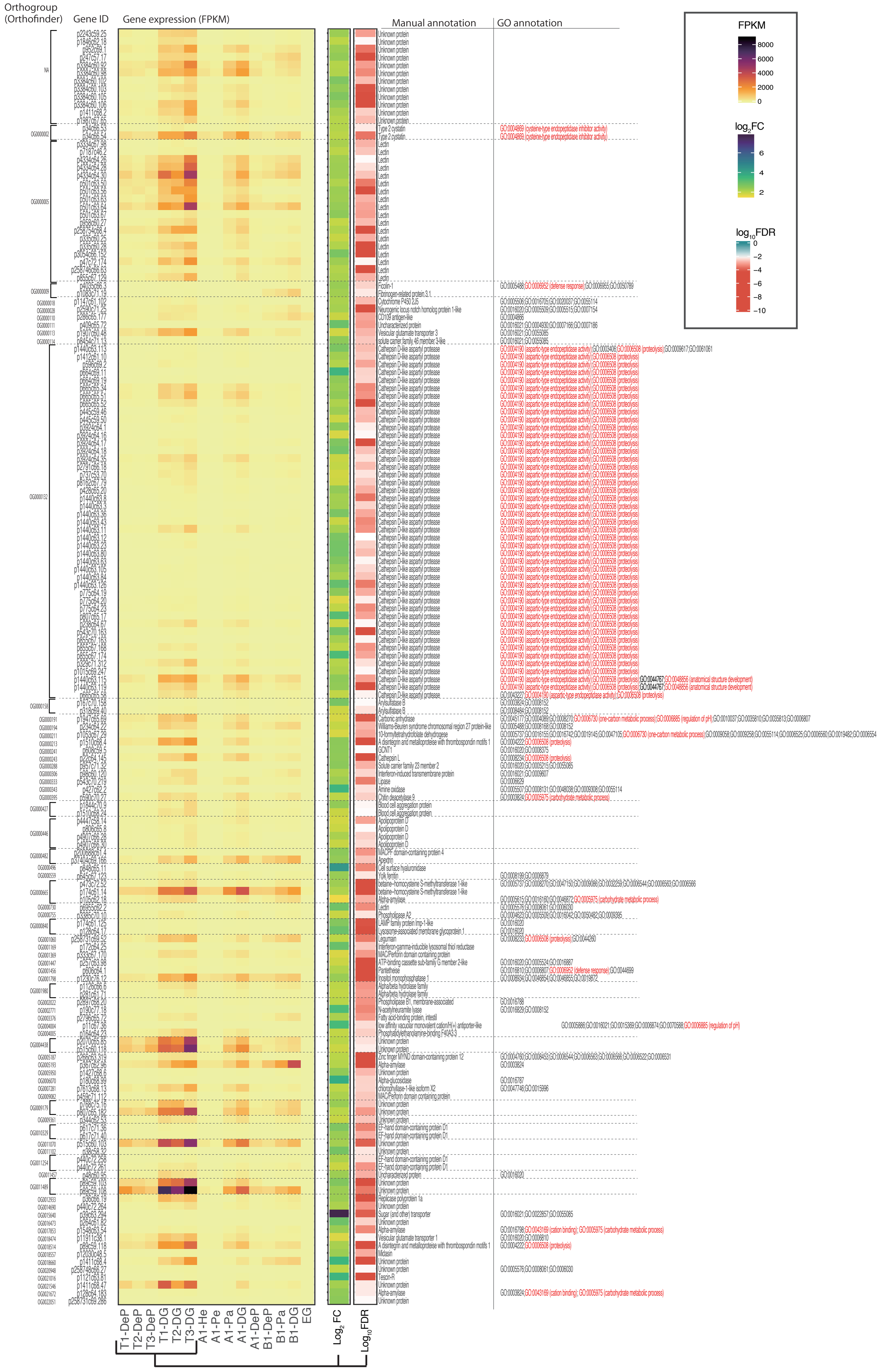

### Figure_5_supple_2.pdf

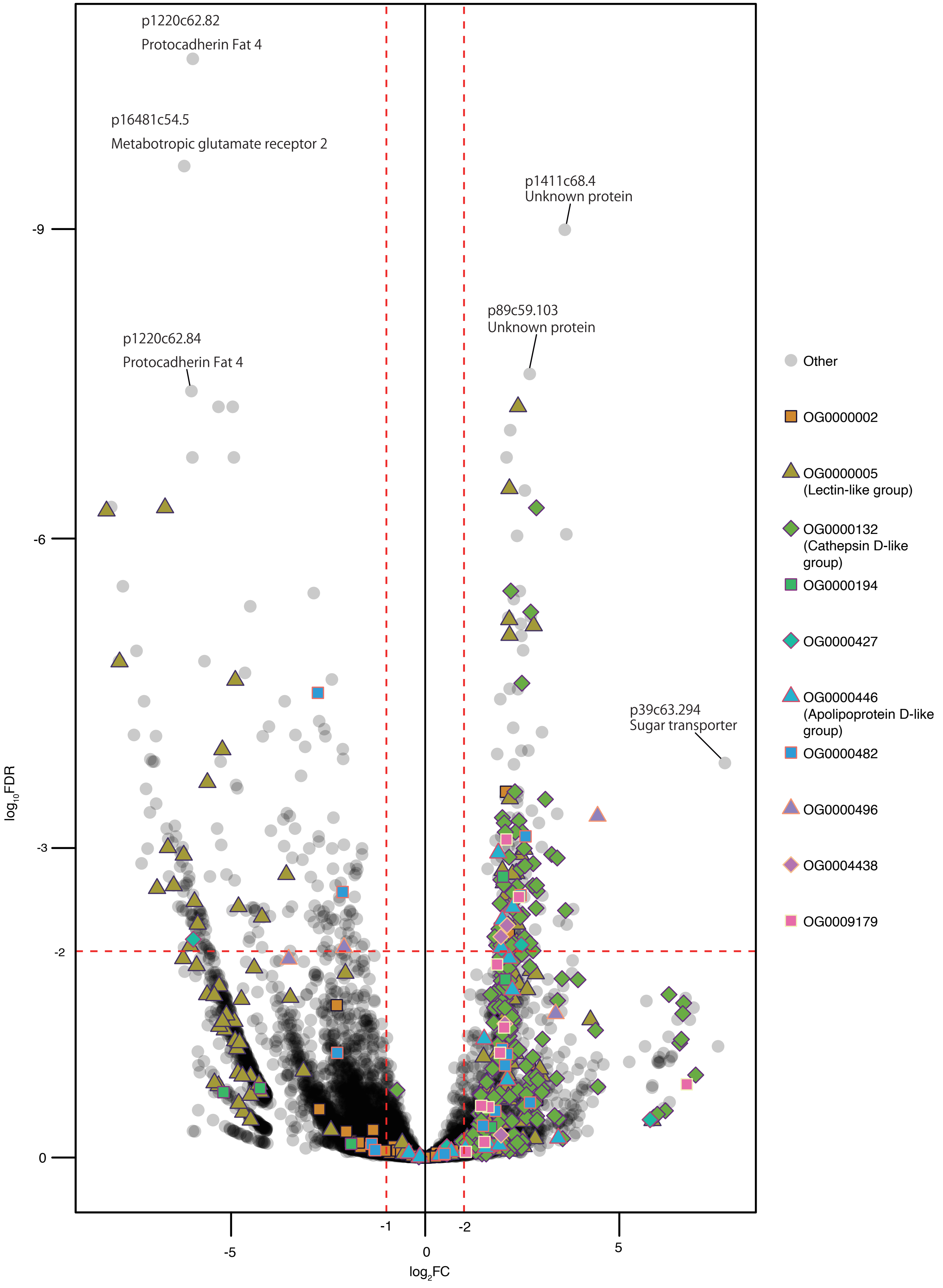

### Figure_5_supple_3.pdf

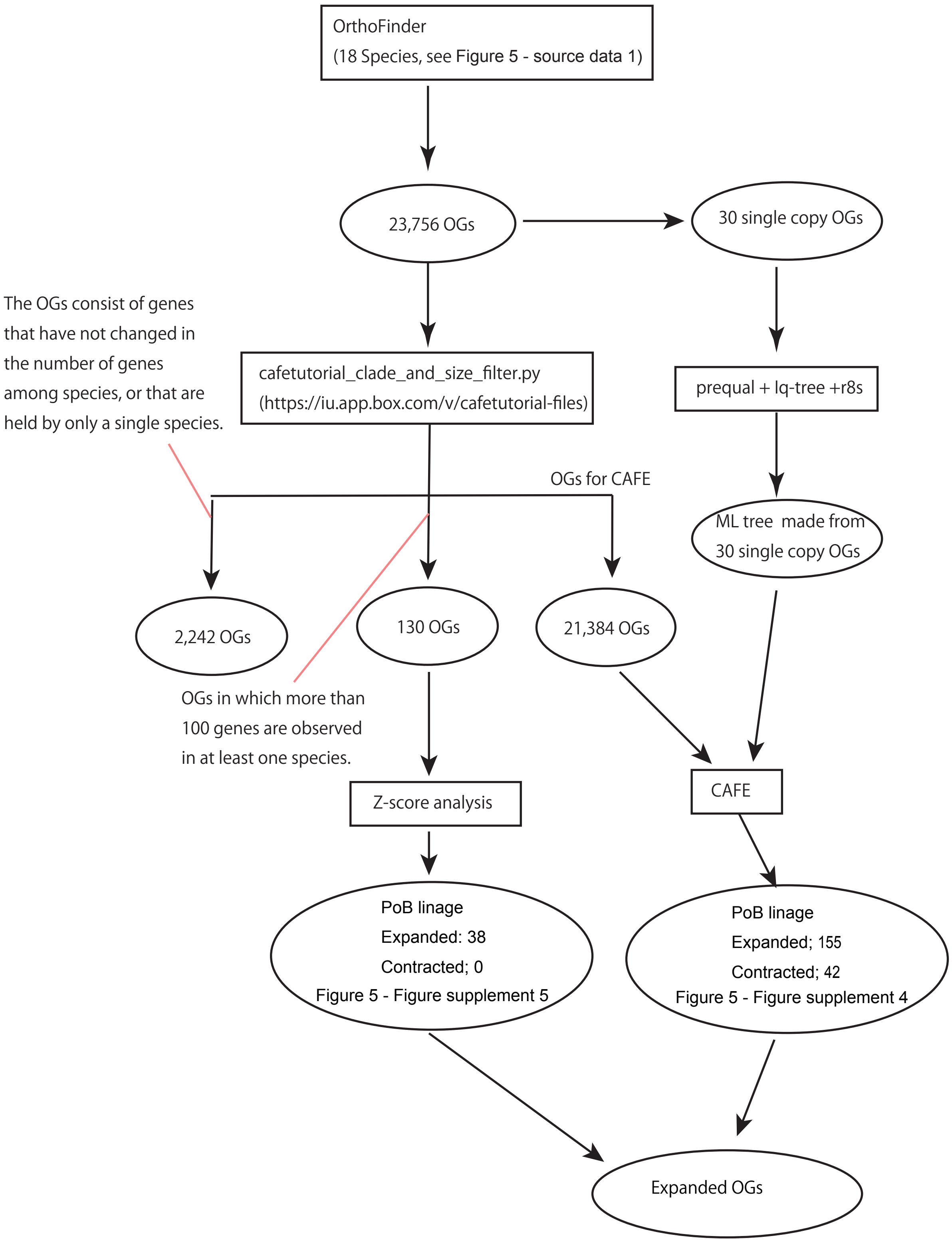

### Figure_5_supple_4.pdf

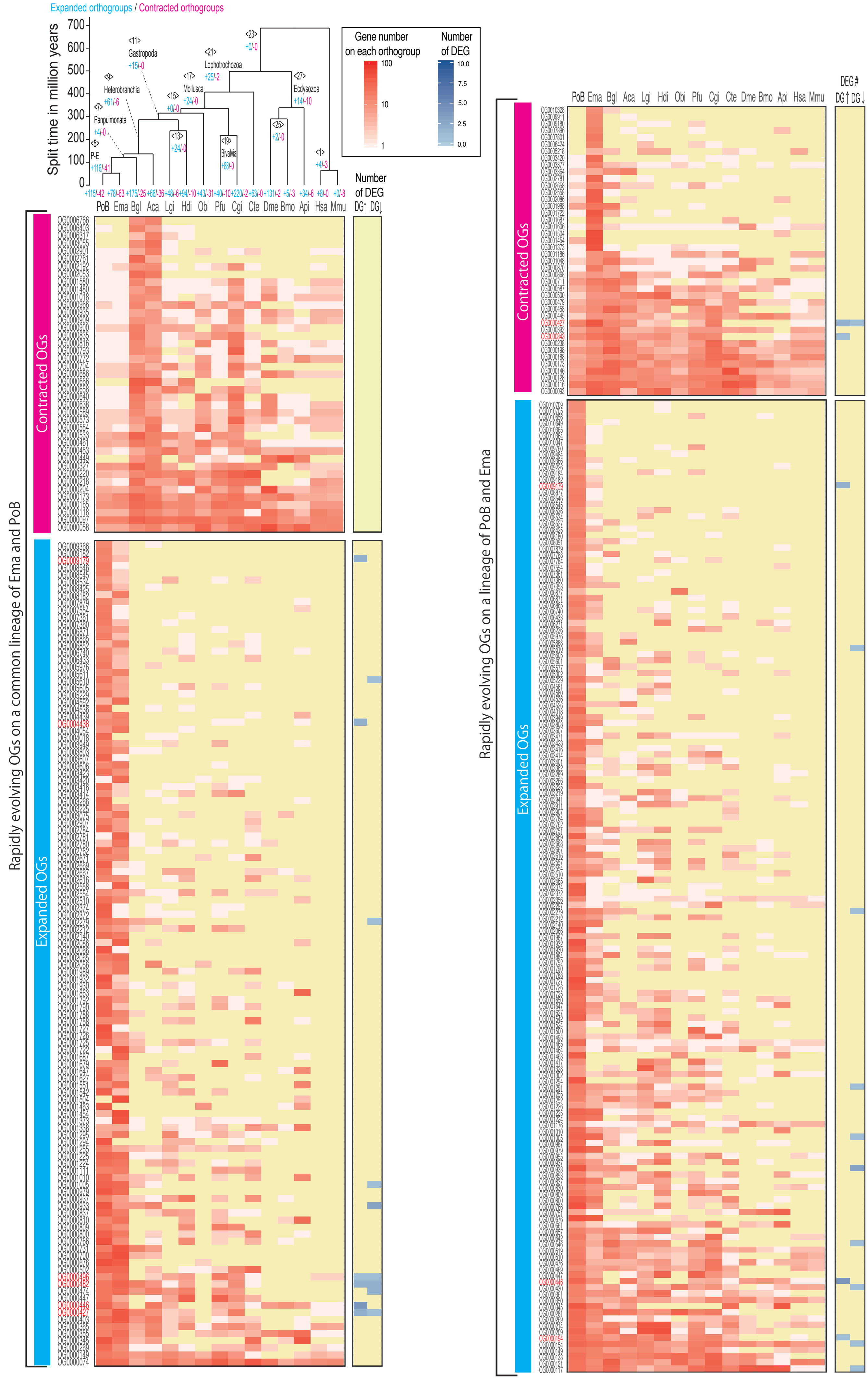

### Figure_5_supple_5.pdf

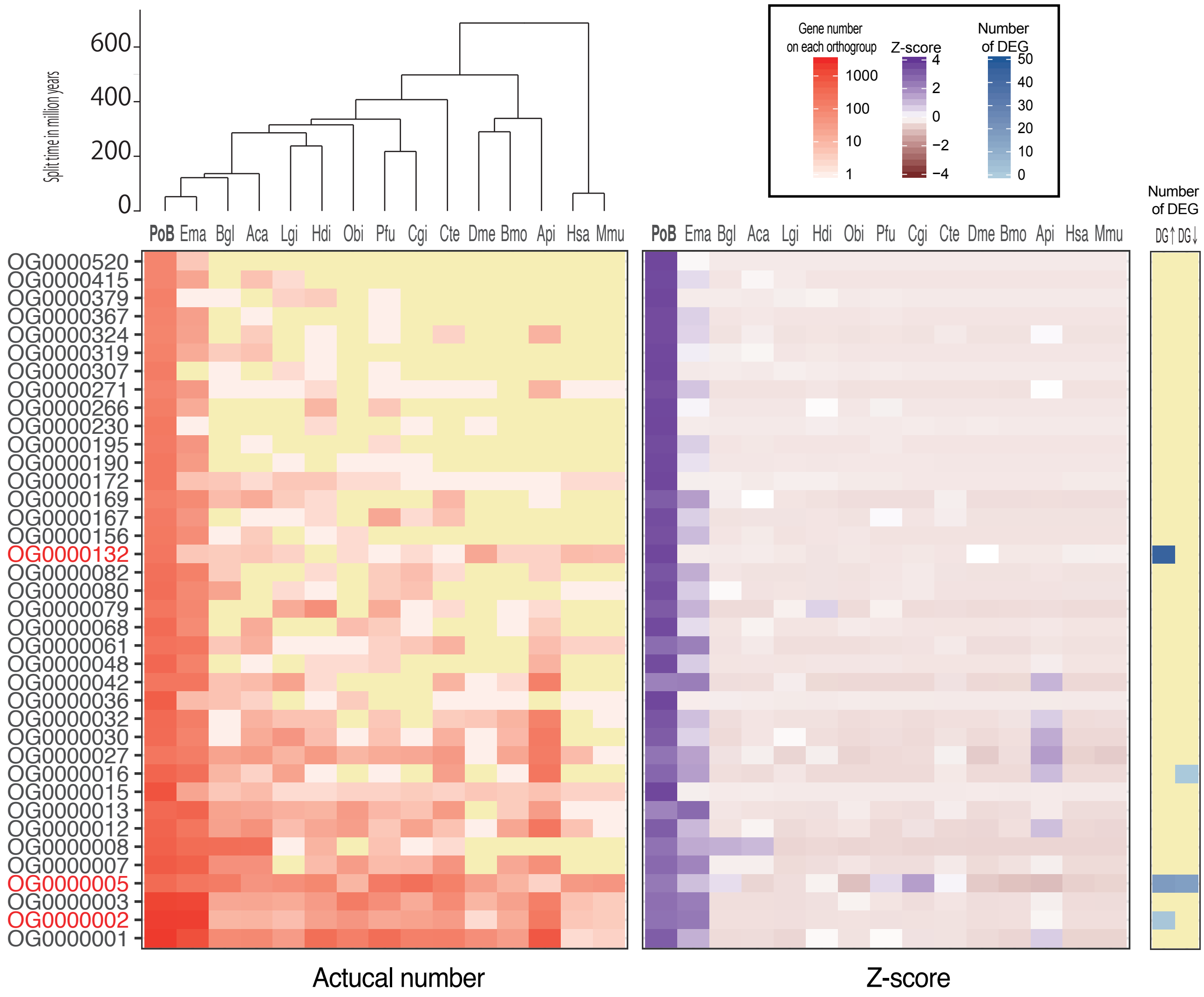

### Figure_6_supple_1.pdf

#### OG0000132: Cathepsin D like

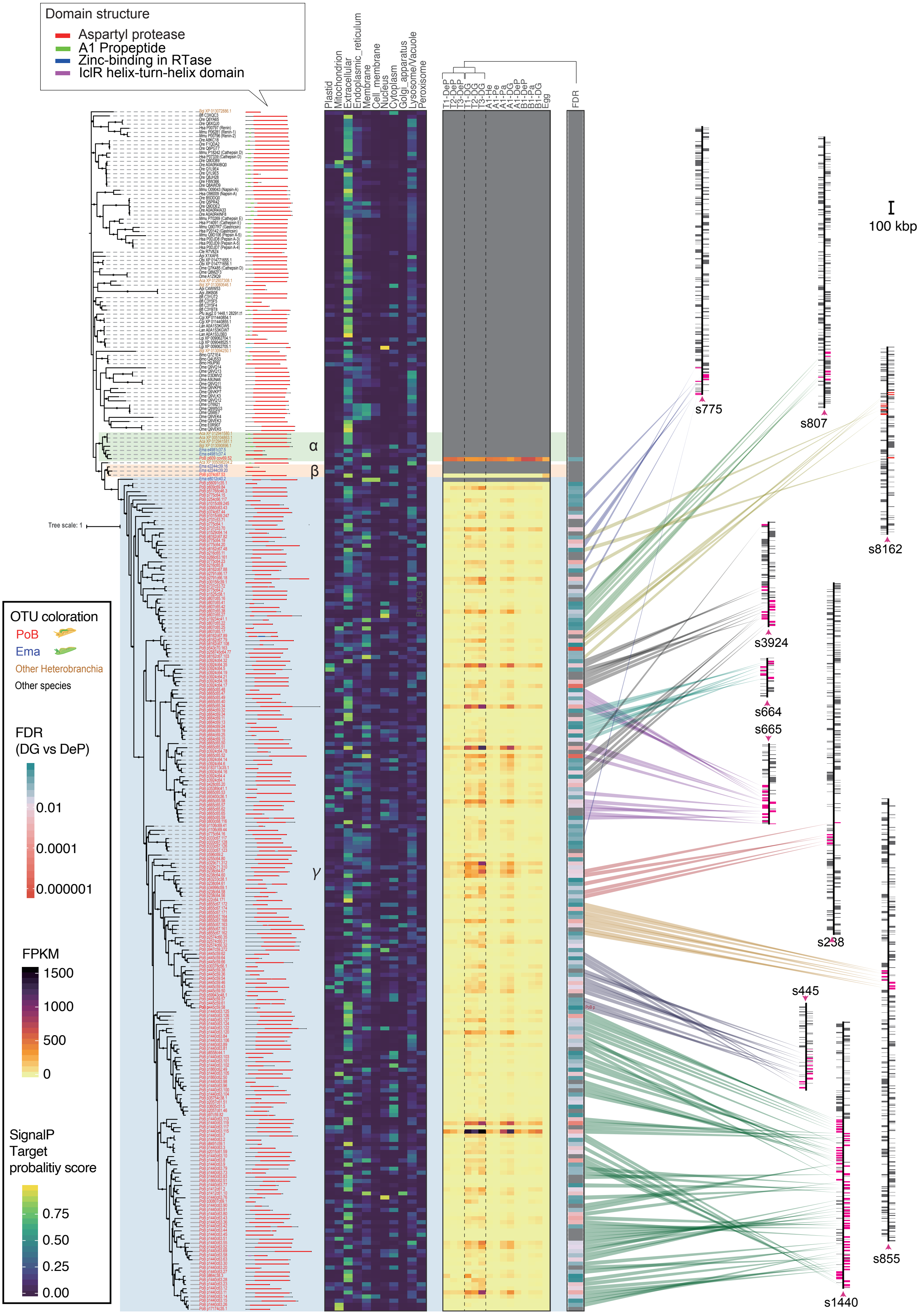

### Figure_6_supple_2.pdf

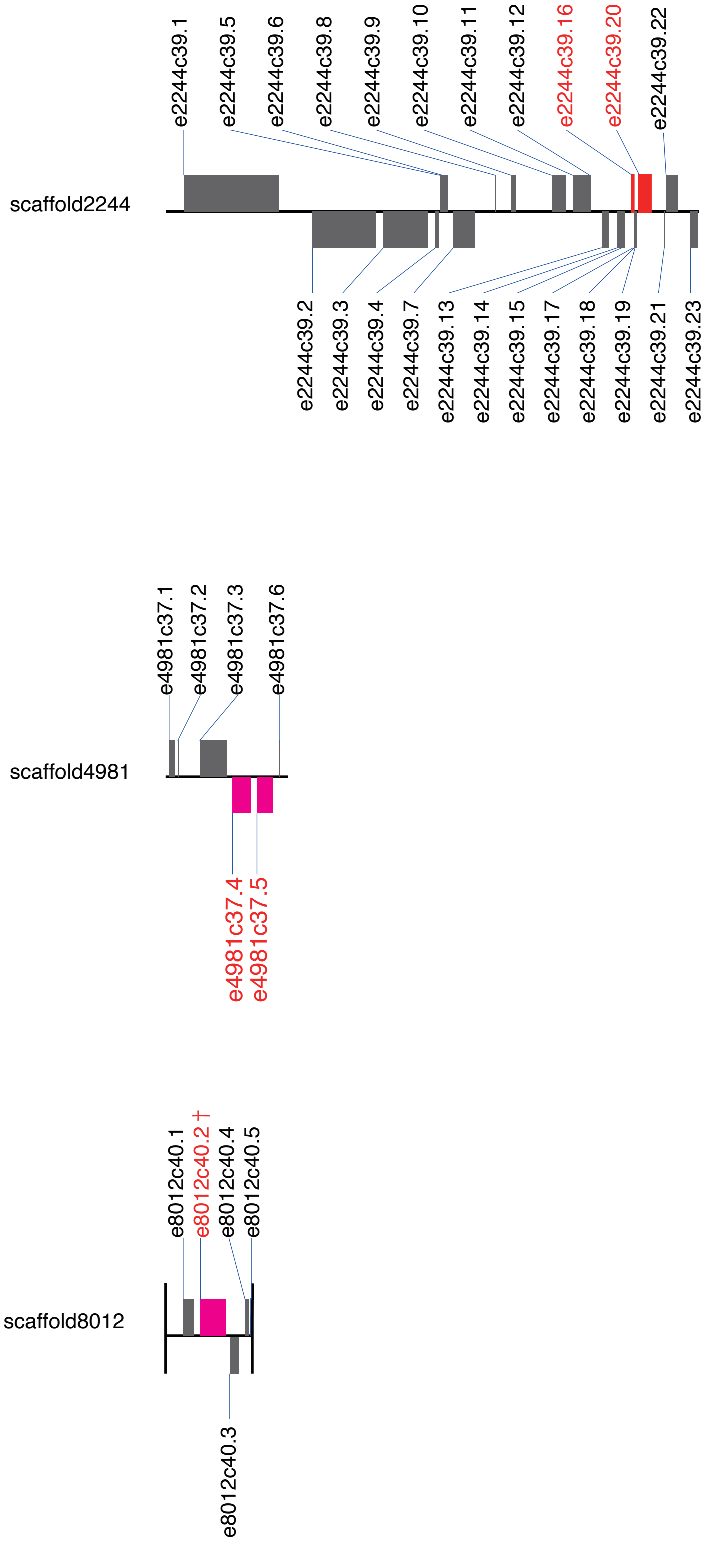

### Figure_6_supple_3.pdf

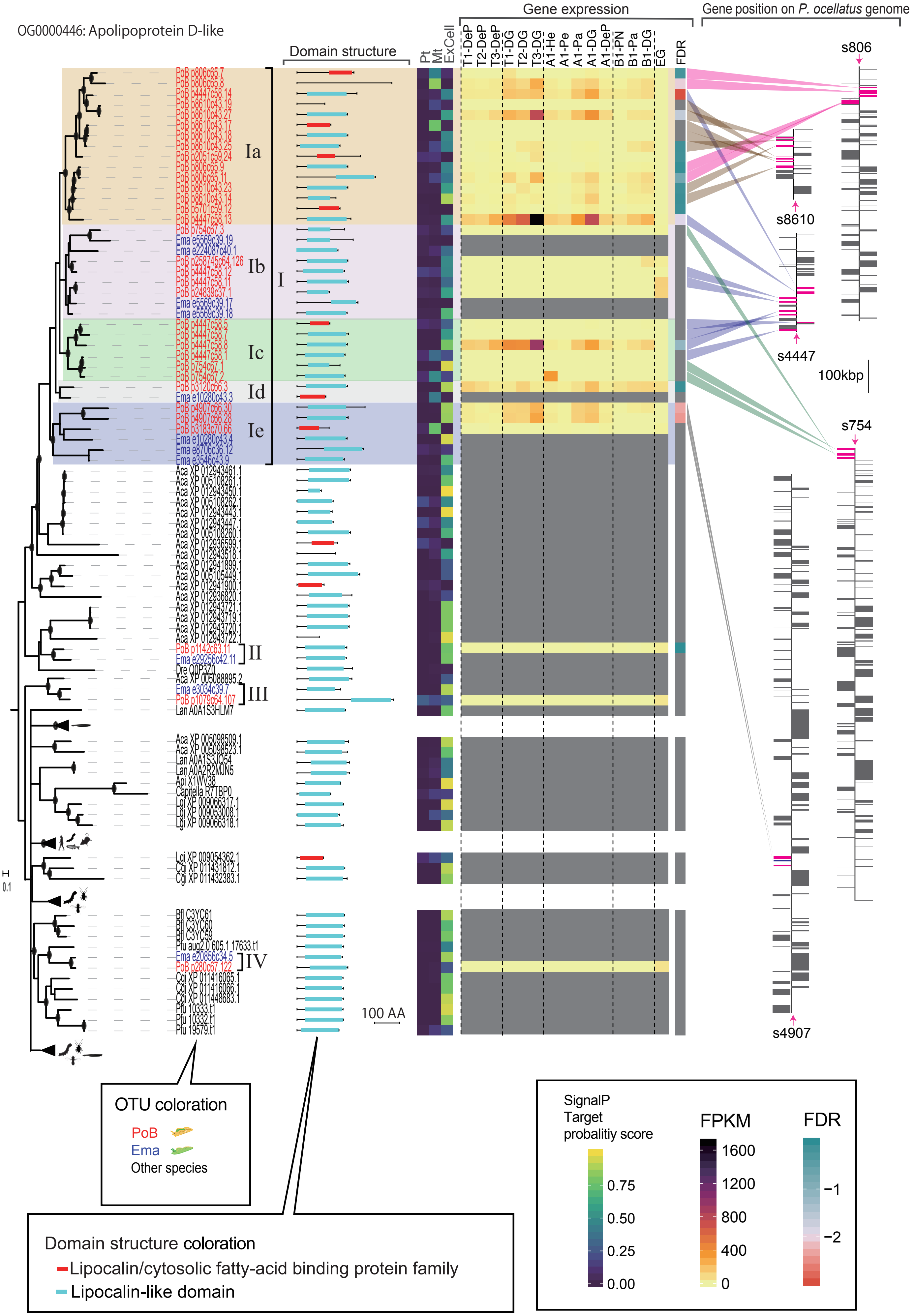

### Figure_6_supple_4.pdf

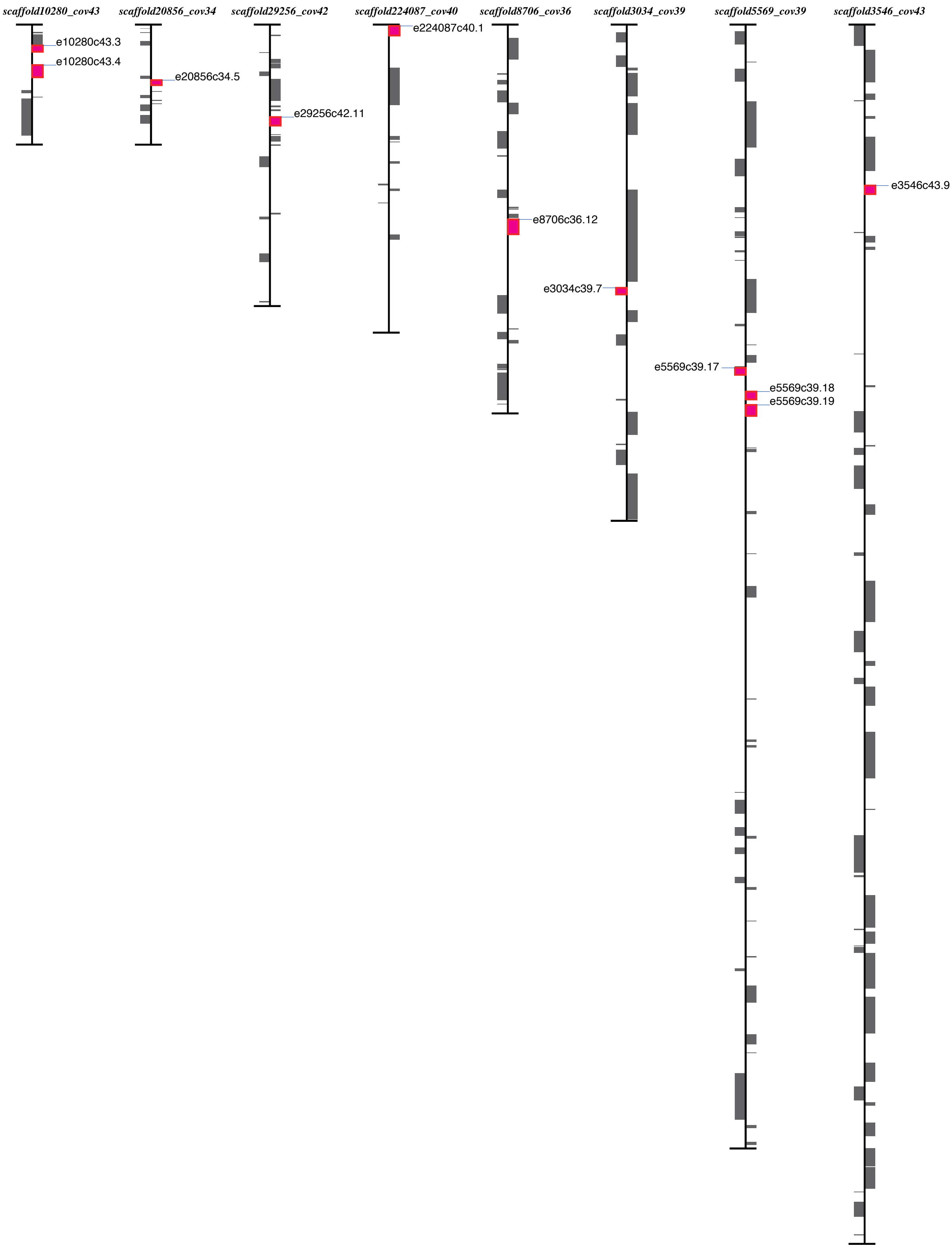
